# Small molecule targeting r(UGGAA)_n_ disrupts RNA foci and alleviates disease phenotype in *Drosophila* model

**DOI:** 10.1101/2020.10.05.323261

**Authors:** Tomonori Shibata, Konami Nagano, Morio Ueyama, Kensuke Ninomiya, Tetsuro Hirose, Yoshitaka Nagai, Kinya Ishikawa, Gota Kawai, Kazuhiko Nakatani

## Abstract

Synthetic small molecules modulating RNA structure and function have therapeutic potential for RNA diseases. Here we report our discovery that naphthyridine carbamate dimer (NCD) targets disease-causing r(UGGAA)_n_ repeat RNAs in spinocerebellar ataxia type 31 (SCA31). Structural analysis of the NCD-UGGAA/UGGAA complex by nuclear magnetic resonance (NMR) spectroscopy clarified the mode of binding that recognizes four guanines in UGGAA/UGGAA pentad by hydrogen bonding with four naphthyridine moieties of two NCD molecules. Biological studies show that NCD disrupts naturally occurring RNA foci built on r(UGGAA)_n_ repeat RNA known as nuclear stress bodies (nSBs) by interfering with RNA-protein interactions resulting in the suppression of nSBs-mediated splicing event. Feeding NCD to larvae of the *Drosophila* model of SCA31 alleviates disease phenotype induced by toxic r(UGGAA)_n_ repeat RNA. These studies demonstrated that small molecules targeting toxic repeat RNAs are a promising chemical tool for studies on repeat expansion diseases.

---

Aberrant expansion of specific repeat sequences in the human genome causes more than 40 neurological disorders called repeat expansion diseases.^1-3^ Expansion of more than 36 CAG repeats in the Huntingtin gene *HTT* and 200 CGG repeats in the FMR1 gene caused Huntington’s disease and Fragile X syndrome, respectively. Extremely long CTG repeats have been identified in the *DMPK* gene of patients of Myotonic Dystrophy type 1 (DM1), although the healthy person has the CTG repeat within the range of 5–38. These diseases have also been classified as trinucleotide repeat diseases because the diseases-causing repeat sequence consisted of three nucleotides. Besides these, diseases caused by the aberrant expansion of tetra-, penta-, and hexanucleotide repeats are also reported.^4-9^

Recent studies on repeat expansion diseases showed that the transcript of aberrantly expanded repeat, thus, long repeat RNA termed as toxic RNAs would be a cause of most repeat expansion diseases.^10–14^ The proposed mechanisms of onset by toxic RNAs are sequestration of RNA-binding proteins (RBPs) in nuclei and unusual translation initiation called repeat-assisted non-AUG (RAN) translation.^15,16^ RNA foci are the aggregates of toxic RNAs and sequester RBPs such as splicing factors in nuclei and dysfunction of these proteins. In contrast, RAN translation produces peptides with the repeated amino acid sequence with the tendency of forming aggregates in the cytoplasm. This gain-of-function of toxic RNAs can account for some characteristics of the pathogenesis of repeat expansion diseases.

Repeat DNA and RNA sequences can fold into hairpin structures involving bulges and mismatches consisting of partially hydrogen-bonded structures, and these structures are potential binding sites for small molecules. Despite the understandings of potential vital roles of small molecules in treating repeat expansion diseases, the exploration of small molecules targeting toxic RNAs remains a challenge. High throughput screening methods including fluorescent indicator-displacement assays,^17–21^ small molecule microarrays,^22,23^ and phenotypic assays^24,25^ have been performed to discover small molecules binding to RNAs. Towards this end, our laboratory has focused on the repositioning of previously developed mismatch binding ligands (MBLs) that target base mismatches in double-stranded DNA (dsDNA).^26-32^ MBLs are designed to bind to the specific mismatches in dsDNA and do not always bind to the mismatches with the same sequence context in double-stranded RNAs (dsRNAs). We anticipated, however, many chances to discover molecules binding to mismatches in dsRNA by screening from our in-house MBL library. We here report that one of MBLs, naphthyridine carbamate dimer (NCD),^29^ binds in high affinity to the 5’-UGGAA-3’/5’-UGGAA-3’ site produced in the hairpin structure of the r(UGGAA)_n_ repeat RNA causing spinocerebellar ataxia type 31 (SCA31). SCA31 is an autosomal dominant spinocerebellar degenerative disorder, and the patients of SCA31 have 2.5–3.8 kbp multiple pentanucleotide repeats containing d(TGGAA)_n_, d(TAGAA)_n_, d(TAAAA)_n_ and d(TAGAATAAAA)_n_ in the genome. Among these repeats, d(TGGAA)_n_ is identified as a pathogenic repeat, as other repeats (i.e., d(TAGAA)_n_, d(TAAAA)_n_ and d(TAGAATAAAA)_n_) is found in unaffected individuals.^33^ Nuclear magnetic resonance (NMR) spectra of the NCD-UGGAA/UGGAA pentad complex reveal the characteristic NCD-bound RNA structure involving the formation of hydrogen bonding pair with the naphthyridine and guanine residue, the zigzag binding orientation of two NCD molecule in the pentad site accompanied with adenine flip out, and the *syn* glycosidic conformation for all naphthyridine-bound guanines. *In vitro* pulldown studies reveal that NCD interfered with the interaction of several RBPs with r(UGGAA)_n_ repeat RNA. NCD treatment resulted in suppression of RNA foci formation in HeLa cells with overexpressed r(UGGAA)_n_ SCA31-related RNA and the inhibition of endogenous nuclear stress bodies (nSBs) assembly upon thermal stress exposure resulting in the disruption of nSBs-dependent splicing regulation. Most significantly, feeding NCD to larvae of *the Drosophila* model of SCA31 results in alleviation of disease phenotype induced by toxic r(UGGAA)_n_ repeat RNA. The observed phenotype changes are most likely due to the interference of the interaction between the r(UGGAA)_n_ repeat RNA and RBPs by NCD as confirmed *in vitro* and cells. Studies described here clearly show that small molecules targeting toxic repeat RNAs have enormous potential to alleviate the toxic RNA function *in vivo*.

Transcription of the (TGGAA)_n_ repeat identified as a SCA31-causing repeat^34^ produced long r(UGGAA)_n_ repeat RNAs,^35^ which formed foci with concomitant co-localization of 43 kDa TAR-DNA binding protein (TDP-43) in addition to more than 80 protein components.^36^ *Drosophila* models of SCA31 expressing long r(UGGAA)_n_ repeat RNA showed degeneration of the eye morphology, life-shortening, locomotor defects, and exhibited similar pathogenesis observed in human SCA31 such as accumulation of r(UGGAA)_n_-containing RNA foci. Coincidentally, the r(UGGAA)_n_ repeat sequence is also found in the HSATIII long noncoding RNAs (lncRNAs), which are transcribed from Satellite III pericentromeric repeat upon thermal stress exposure.^37^ HSATIII lncRNAs also sequestrate multiple RBPs to form RNA foci called nSBs in the nucleus. The formation of nSBs results in the suppression of pre-mRNA splicing (or promote intron retention) of ∼500 introns during recovery from thermal stress.^38^ Overexpression of motor neuron disease-associated RBPs such as TDP-43 and fused in sarcoma (FUS) in *Drosophila* models of SCA31 mitigated toxicity of r(UGGAA)_n_ repeat RNA accompanied with the dispersion of RNA foci.^36^ These results suggested that binding of these RBPs to r(UGGAA)_n_ repeat RNA induced the release of the sequestered proteins from the RNA and, more importantly, suggest the possibility of small molecules in reducing the RNA toxicity by their binding to r(UGGAA)_n_ repeat RNA. This hypothesis is relevant to those proposed for small molecules targeting disease-causing toxic RNAs including r(CUG)_n_ in DM1,^39–41^ r(CCUG)_n_ in DM2,^42,43^ r(CGG)_n_ in fragile X-associated tremor/ataxia syndrome,^44,45^ r(GGGGCC)_n_ in ALS/FTD,^46,47^ and r(AUUCU)_n_ in SCA10.^48^ We, therefore, started to screen our in-house chemical library to find molecules binding to r(UGGAA)_n_ repeat RNA.

## Results

### Screening of small molecules binding to r(UGGAA)_n_ repeat RNAs from the in-house chemical library

To explore molecules binding to r(UGGAA)_n_ repeat RNAs, we have screened our in-house MBL library by surface plasmon resonance (SPR) assay, where three repeat RNAs r(UGGAA)_9_, r(UAGAA)_9_, and r(UAAAA)_9_ were immobilized on its surface. Non-pathogenic r(UAGAA)_9_ and r(UAAAA)_9_ repeats were used as control RNAs. Most disease-causing repeat RNAs responsible for RNA-mediated neurodegeneration can form hairpin structures involving internal loops consisting of multiple mismatched base pairs.^49^ The r(UGGAA)_n_ repeat RNA may fold into the hairpin structures, which include the UGGAA/UGGAA pentad containing the 5’-GGA-3’/3’-AGG-5’ internal loop (Fig. 1a). From our previous studies, the guanine-rich internal loop could be good target motifs for MBLs consisting of two *N*-acyl-2-amino-1,8-naphthyridine heterocycles, of which hydrogen bonding surface is complementary to that of guanine base (Fig. 1b). Among 20 compounds we investigate (LC-1∼LC-20) (Supplementary Fig. 1), four compounds (LC-5, 8, 9, and 11) showed the marked increase in the SPR signal to the r(UGGAA)_9_-immobilized surface (Fig. 1c and Supplementary Fig. 2) and the band shift of r(UGGAA)_9_ by gel electrophoretic mobility shift assay (EMSA) (Fig. 1d) with concentration-dependent manner (Supplementary Fig. 3). Structural comparison of 4 compounds readily suggested that LC-9, which is called NCD showing high affinity to the G-rich DNAs^29,30^ would be the minimum structural element for the binding to r(UGGAA)_9_. The other three compounds were either dimeric form or a very close derivative of NCD.

**Fig. 1.**
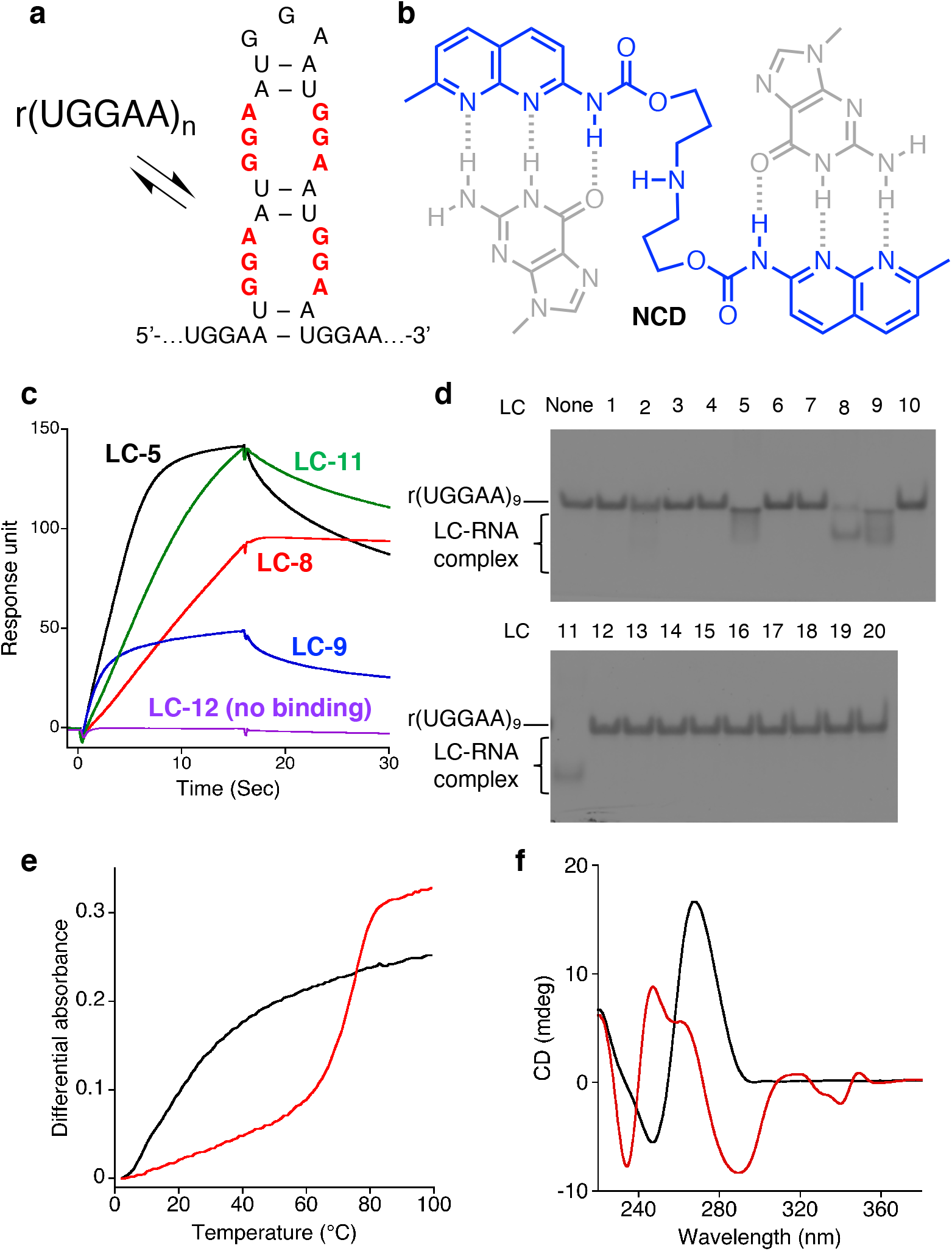
Screening of r(UGGAA)_n_-binding small molecules from an in-house chemical library. **a**, Possible hairpin structures in r(UGGAA)_n_ containing internal loop with three consecutive G-A, G-G and A-G mismatch base pairs (shown in bold). **b**, Chemical structure of NCD (shown in blue) and schematic illustration of hydrogen bonding to guanines (shown in gray). **c**, Representative SPR data for the binding of LC-5, 8, 9, and 11, and for the non-binding of LC-12 to r(UGGAA)_9_-immobilized surface. **d**, Native PAGE analysis of interactions between LC-1–20 and r(UGGAA)_9_. **e**, UV melting profiles and **f**, CD spectra of r(UGGAA)_9_ in the absence (black) or presence (red) of NCD.

To gain insights into the NCD-binding to r(UGGAA)_9_, we carried out structure-binding relationship studies on NCD derivatives by EMSA (Supplementary Fig. 4). Neither compounds having different linker structure connecting two naphthyridine heterocycles (LC-2 and 3) nor the molecule having only one naphthyridine heterocycle showed the band shift for the r(UGGAA)_9_. These results confirmed that NCD is the indispensable structural element for the binding to r(UGGAA)_9_. UV melting profiles measured for the r(UGGAA)_9_ in the presence of NCD (20 μM) showed the melting temperature (*T*_m_) of 73.7 °C (Fig. 1e), whereas the RNA itself did not show *T*_*m*_. Such a markedly high *T*_*m*_ was not observed for the r(UAGAA)_9_ in the presence of NCD, and the r(UAAAA)_9_ did not show any increase in *T*_*m*_ by NCD (Supplementary Fig. 5). Substantial changes in the circular dichroism (CD) spectra of the r(UGGAA)_9_ observed upon NCD-binding (Fig. 1f) suggested the formation of NCD-bound complex with marked structural changes.

### Sequence selectivity in the binding of NCD to mutated UGGAA-UGGAA pentad

To know if NCD-binding to the r(UGGAA)_9_ involved the binding to the UGGAA/UGGAA pentad in the secondary hairpin structure, *T*_m_s of the RNA duplexes containing the U**RRA**A/U**RRA**A pentad, where **R** is guanine (G) or adenine (A), were measured in the absence and presence of NCD (Fig. 2a and Supplementary Fig. 6). A marked increase in the *T*_*m*_ by 21.5 °C was observed for the duplex containing the UGGAA/UGGAA pentad (5’-**GGA**-3’/3’-**AGG**-5’) in the presence of NCD (red bar), whereas no increase in the *T*_m_ was observed with quinoline carbamate dimer (QCD) (blue bar). QCD did not show the band shift for r(UGGAA)_9_ in EMSA (Supplementary Fig. 4). Replacing one guanine in 5’-**GGA**-3’/3’-**AGG**-5’ with adenine to 5’-**GGA**-3’/3’-**AG<u>A</u>**-5’ (underlined) resulted in a modest *T*_m_ increase (5.2 °C) in the presence of NCD. Further substitution of the guanine in 5’-**GGA**-3’/3’-**AGA**-5’ with adenine to 5’-**<u>A</u>GA**-3’/3’-**AGA**-5’ and 5’-**GGA**-3’/3’-**A<u>A</u>A**-5’ diminished the effect of NCD on the increase in *T*_m_. Substantial changes on CD spectra of the duplex containing the UGGAA/UGGAA pentad were also observed in the presence of NCD but not QCD (Fig. 2b). These results suggested that NCD binding to the r(UGGAA)_9_ likely involved the binding to the UGGAA/UGGAA pentad. With the 29mer hairpin RNA containing the UGGAA/UGGAA pentad (SCA31 hpRNA; Supplementary Fig. 7), the stoichiometry of NCD binding to the UGGAA/UGGAA pentad was determined to be 2:1 by electrospray ionization time-of-flight mass spectrometry (ESI-TOF MS) analysis (Fig. 2c and Supplementary Fig. 8). The ions corresponding to the 1:1 complex was not detected under the conditions, suggesting that binding of two NCD molecules to the UGGAA/UGGAA pentad would be necessary to stabilize the NCD-SCA31 hpRNA complex. Accurate thermodynamic parameters for the binding of NCD to the UGGAA/UGGAA pentad were determined by isothermal titration calorimetry (ITC) to give *ΔH* –10.6 kcal·mol^−1^, *ΔS* –1.3 cal·mol^−1^·K^−1^, *and ΔG* –10.2 kcal·mol^−1^ (at 25 °C) leading to an apparent dissociation constant *K*_*d(app)*_ of 32 nM (Fig. 2d).

**Fig. 2.**
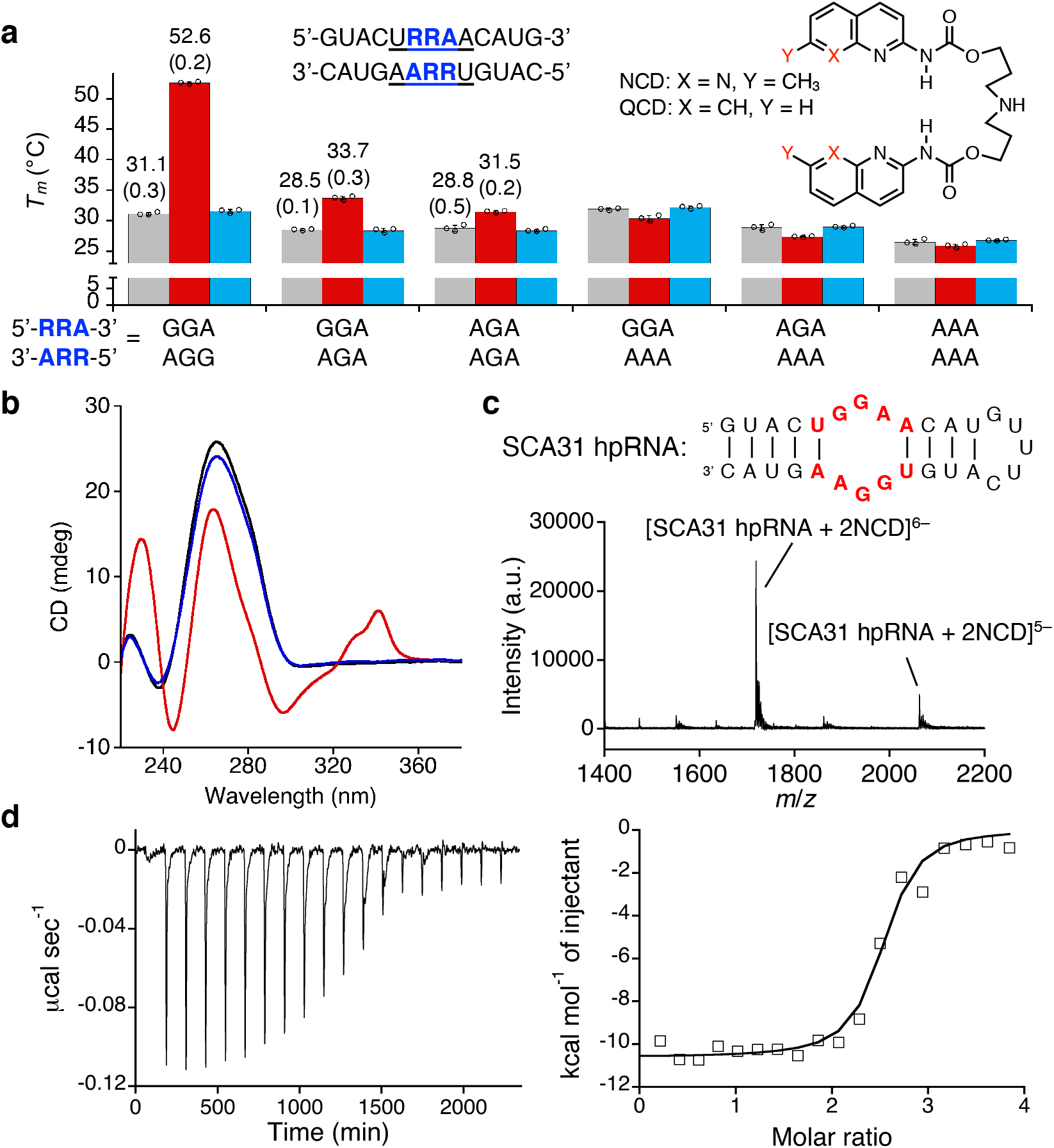
Binding of NCD to UGGAA/UGGAA pentad-containing RNA. **a**, Melting temperatures of RNA duplexes containing URRAA/URRAA pentad in the absence (gray) and the presence of NCD (red) and QCD (blue). The data in *T*_m_ measurements are presented as the mean ± SD (*n* = 3). **b**, CD spectra of RNA duplexes containing UGGAA/UGGAA pentad (black) in the presence of NCD (red) and QCD (blue). **c**, ESI-TOF-MS spectra of UGGAA/UGGAA pentad-containing hairpin RNA (10 μM) in the presence of 40 μM NCD. **d**, ITC measurements for the binding of NCD to UGGAA/UGGAA pentad-containing hairpin RNA.

### Structural analysis of NCD-UGGAA/UGGAA pentad complex by NMR

Interactions of NCD to the SCA31 hpRNA (Fig. 3a) were analyzed by NMR spectroscopy. Fig. 3b showed the changes in NMR signals of exchangeable protons upon the addition of NCD, indicating the structural change of SCA31 hpRNA. Signals that appeared around 10-12 ppm were due to the formation of four naphthyridine-guanine hydrogen-bonded pairs, which were confirmed by the NOE, as shown in Fig. 3c. It is noted that signals of guanosine residues of the UGGAA/UGGAA pentad could not be observed without NCD, likely due to the exchange broadening, suggesting that the UGGAA/UGGAA region is structurally flexible.

**Fig. 3.**
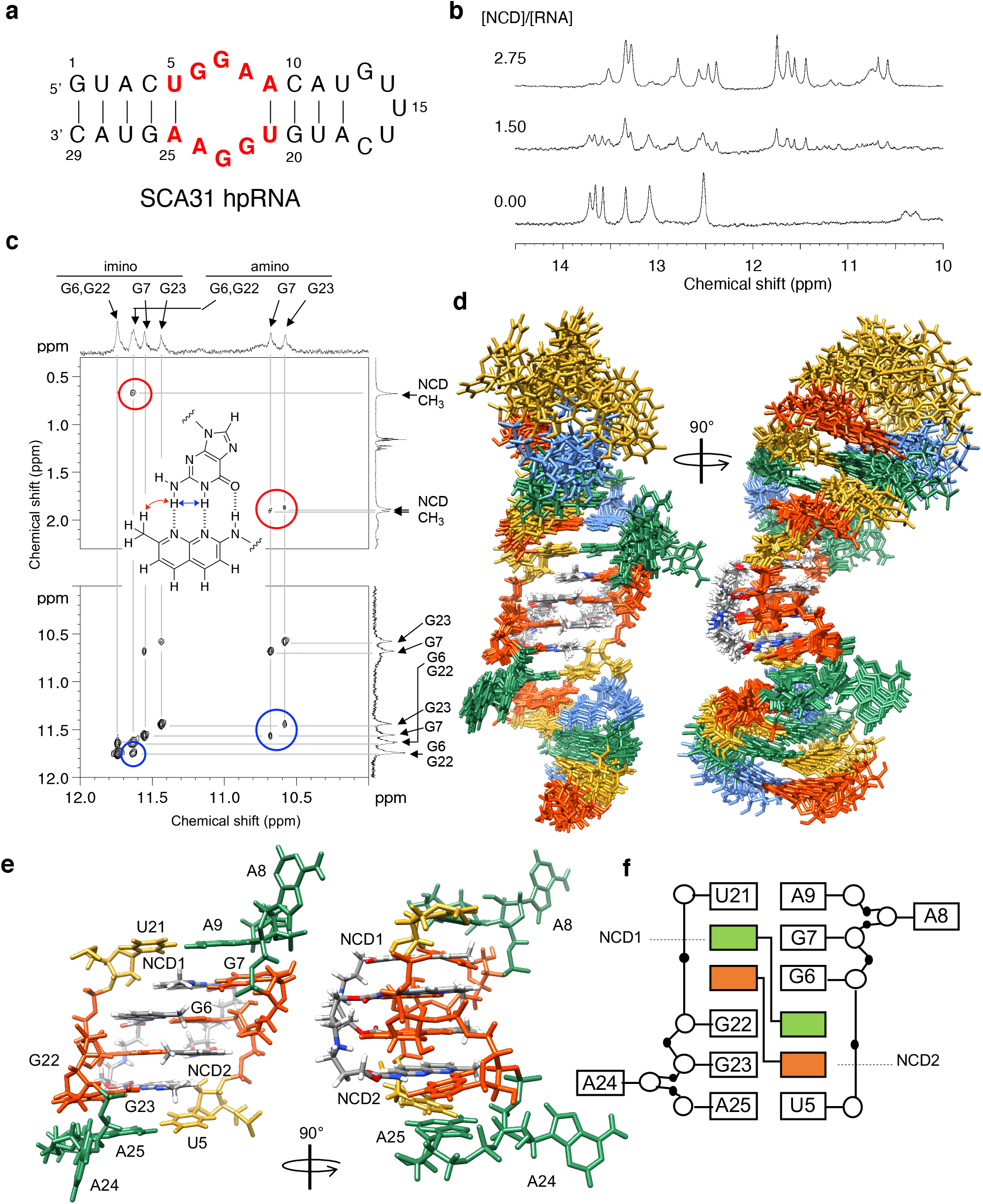
NMR structural analysis of NCD-UGGAA-UGGAA pentad complex. **a**, SCA31 hpRNA used for the NMR analysis. **b**, Changes in imino proton spectra of SCA31 hpRNA upon addition of NCD. **c**, NOESY spectra of NCD-SCA31 hpRNA complex. Correlations between amino hydrogen and methyl group (red circle) and amino and imino hydrogens (blue circle) were shown with the assignment of hydrogens. **d**, Ensemble of the ten lowest total energy structures of the NCD-SCA31 hpRNA complex. **e**, Close-up structures and **f**, structure pattern diagram of the UGGAA-UGGAA pentad bound with two NCD molecules.

Both exchangeable and non-exchangeable protons of the SCA31 hpRNA-NCD complex were analyzed, and several intermolecular NOEs were found, as shown in Fig. 3c and Supplementary Fig. 9, for example. NMR signals indicated the symmetry in the structure around the UGGAA/UGGAA pentad because the signal of G6 is overlapped with that of G22, and signals of G7 and G23 are close to each other. Signals of four methyl groups in two NCD molecules were observed at 0.7 and around 1.9 ppm, suggesting that two NCD molecules in UGGAA/UGGAA pentad adopt a symmetric structure. The superposition of the ten lowest energy structures was shown in Fig. 3d and the structural statistics were summarized in Supplementary Table 1. Structures of the UGGAA/UGGAA pentad with two NCD molecules were well defined, and the minimized average structure of the region was shown in Fig. 3e and 3f. The four guanosine residues bound by naphthyridine were found to be in *syn* conformation regarding the *N*-glycosidic bond; therefore, NCD bound to the UGGAA/UGGAA pentad from the minor groove. The four naphthyridine-guanine base pairs are stacked between the two A-U base pairs, and the two A residues (A8 and A24) are flipped out toward the major groove. The linker of naphthyridine moieties is located in the minor groove connecting G6 and G23 as well as G22 to G7. It is noted that only the current combination of naphthyridine-guanine pairs provided us acceptable converged structures. Four methyl groups of two NCD molecules are located in line to form a hydrophobic core in the major groove. These structural features of cross-connections and hydrophobic core may contribute to the stabilization of the complex. Two of four methyl groups in NCD were observed in the high field (0.7 ppm), whereas the other two methyl groups were observed at around 1.9 ppm. Naphthyridine rings opposite to G7 and G23 probably caused the ring current shifts observed for the two methyl groups of naphthyridine opposite to G6 and G22, respectively. It is important to note that the methyl group of free NCD was observed at 2.0 ppm. We also analyzed the overall quality of the structure by MolProbity^50^ (Supplementary Table 2).

### Inhibitory effect of NCD on the interaction of r(UGGAA)_n_ with RBPs

The high-affinity binding of NCD to the r(UGGAA)_n_ repeat RNA encouraged us to assess the effect of NCD on the interaction between r(UGGAA)_n_ repeat RNA and RBPs. To investigate the effect of NCD on the RNA-protein interactions, we performed *an in vitro* pulldown assay using biotinylated (UGGAA)_20_ with HeLa nuclear extract in the absence and presence of NCD (Fig. 4a). Among RBPs we have tested, TDP-43, heterogeneous nuclear ribonucleoprotein M (HNRNPM), serine and arginine rich splicing factor 9 (SRSF9), and FUS were co-precipitated with the r(UGGAA)_20_, but not with the antisense sequence r(UUCCA)_20_ (Fig. 4b), indicating the sequence-specific binding of these proteins with r(UGGAA)_20_ RNA. In the presence of NCD, we could confirm the inhibitory activity on the co-precipitation of TDP-43, HNRNPM and SRSF9 with the (UGGAA)_20_ (Fig. 4c,d), but moderate or marginal effects on the co-precipitation of FUS, splicing factor proline and glutamine rich (SFPQ), and ALYREF (Fig. 4b). These results suggested that NCD could interfere with the interactions between r(UGGAA)_n_ repeat RNA and TDP-43, HNRNPM, SRSF9, and possibly with other RBPs interacting with SCA31 RNA.

**Fig. 4.**
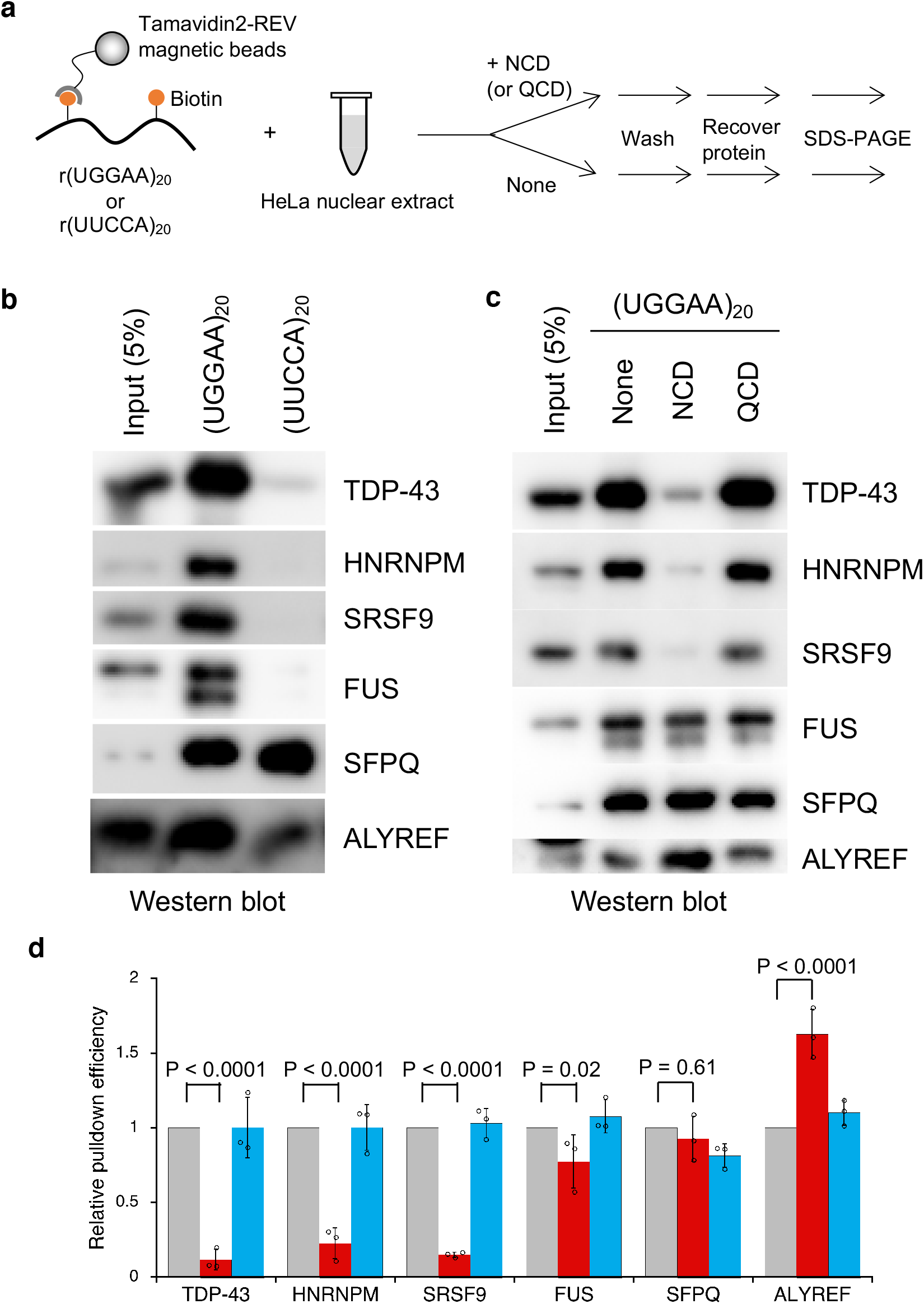
Inhibition of interaction between r(UGGAA)_n_ and RBPs by NCD. **a**, Workflow of *In vitro* pulldown assay. **b**, *In vitro* pulldown assay of RBPs using biotinylated r(UGGAA)_20_ or antisense r(UUCCA)_20_. The co-precipitated proteins were detected by western blotting using specific antibodies. **c**, *In vitro* pulldown assay using biotinylated r(UGGAA)_20_ in the absence and the presence of 2 μM NCD and QCD. **d**, Relative pulldown efficiencies of each protein in the absence (gray) and the presence of NCD (red) and QCD (blue). Data are shown as the mean ± SD (n = 3); (Dunnett’s multiple comparison test).

### Inhibition of assembly of RNA foci and nSBs in HeLa cells

The inhibitory activities of NCD on RNA-protein interactions prompted us to assess the effects of NCD on the assembly of RNA foci and nSBs in HeLa cells. We first analyzed HeLa cells expressing exogenous r(UGGAA)_76_ to examine if NCD disrupts the formation of the RNA foci containing r(UGGAA)_n_ repeat RNA by fluorescence *in situ* hybridization (FISH). In the HeLa cells transfected with the plasmid for expressing r(UGGAA)_76_, the accumulation of RNA foci was observed after incubation for 24 h in the nuclei as confirmed by DAPI staining (Fig. 5a; left). In contrast, HeLa cells pre-incubated with NCD showed significantly reduced formation of RNA foci (Fig. 5a,b; middle). The decrease of RNA foci was not observed by pre-incubation with QCD (Fig. 5a,b; right). It is important to note that NCD did not interfere with the interaction of the FISH probe with r(UGGAA)_n_ repeat RNA, the proliferation of HeLa cells, and the expression of r(UGGAA)_76_ under the conditions (Supplementary Fig. 10). We also observed that NCD interfered with colocalization of TDP-43 with r(UGGAA)_76_, confirming that TDP-43 was released from foci (Supplementary Fig. 11).

**Fig. 5.**
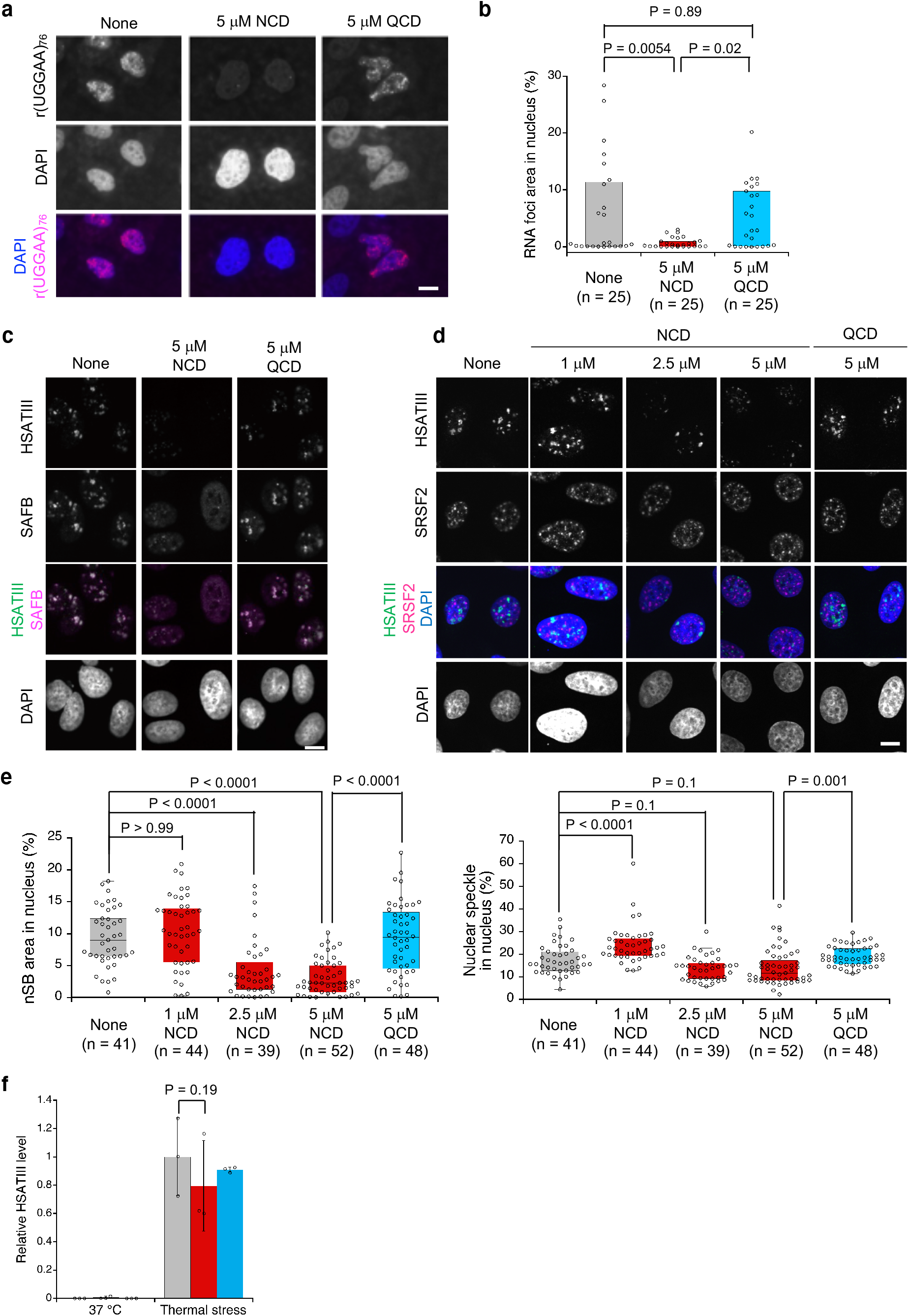
The effect of NCD on RNA foci and nSBs assembly. **a**, RNA FISH images of cells expressing r(UGGAA)_76_ in the absence (left) and presence of NCD (middle) and QCD (right). Ligand concentration was 5 μM. Scale bar, 10 μm. **b**, Box plot of total area of r(UGGAA)_76_ RNA foci in each nucleus. p-value (Kruskal-Wallis test, followed by Dunn’s multiple comparison test) was shown above. **c**,**d** RNA FISH and IF images of HeLa cells after thermal stress exposure in the absence and the presence of NCD (1, 2.5, and 5 μM) or QCD (5 μM) stained by HSATIII-FISH and IF using an anti-SAFB (**c**) and anti-SRSF2 (**d**) antibodies. Scale bar: 10 μm. **e**, Box plot of total area of nSBs (left) and nuclear speckles (right) in each nucleus. p-value (Kruskal-Wallis test, followed by Dunn’s multiple comparison test) was shown above. **f**, RT-qPCR analysis of HSATIII RNA level. HeLa cells were cultured at 37 °C for 3 h (left; 37 °C) or at 42 °C for 2 h followed by recovery at 37 °C for 1 h (right; thermal stress) in the absence (gray) and presence of 5 μM NCD (red) or QCD (blue). Data are shown as the mean ± SD (n = 3); (Dunnett’s multiple comparison test).

HSATIII lncRNAs are induced by thermal stress and sequester various RBPs, including TDP-43, HNRNPM, SRSF9, and FUS, leading to the formation of nSBs.^38,51,52^ The effects of NCD on the inhibition of RNA foci assembly raised the intriguing possibility that NCD may inhibit the nSB formation and function in thermal stress-exposed cells by interfering with the sequestration of nSB proteins. We analyzed the effect of NCD on the assembly of nSBs by RNA FISH and immunofluorescence (IF). In the control and QCD-treated HeLa cells, nSBs were normally formed upon thermal stress exposure at 42 °C for 2 h followed by recovery at 37 °C for 1 h and scaffold attachment factor B (SAFB) (Fig. 5c) and SRSF9 (Supplementary Fig. 12) were detected in nSBs. In contrast, in the presence of NCD, nSBs were decreased in a concentration-dependent manner (Fig. 5d), and SAFB and SRSF9 were diffused throughout the nucleoplasm (Fig. 5c and Supplementary Fig. 12). As compared with nSBs, the NCD treatment gave only a weak effect on another nuclear body, such as nuclear speckle, including SRSF2 (Fig. 5d,e). NCD may weakly bind to the G-rich SRSF2-binding site of GGNG to induce weak effect on nuclear speckle. The reverse transcription-quantitative PCR (RT-qPCR) analysis confirmed that the amount of HSATIII lncRNAs in NCD-treated cells is comparable with those in the control cells (Fig. 5f), suggesting the disruption of nSBs is not due to the degradation of HSATIII. These observations suggested that NCD interfered with the assembly of the r(UGGAA)_n_ repeat RNA-associated RNA foci and nSBs in HeLa cells.

### Effects of NCD on nSB-dependent intron retention

nSBs serve as a part of auto-regulation mechanisms of phosphorylation of SRSFs and splicing of *CLK1* pre-mRNA during thermal stress and subsequent stress recovery (Fig. 6a).^38^ Introns 3 and 4 of *CLK1* pre-mRNA are retained in normal conditions (e.g. 37 °C) and excised upon thermal stress exposure (e.g. 42 °C) which is promoted by thermal stress-dependent de-phosphorylation of SRSFs. During stress recovery, the intron-retaining *CLK1* pre-mRNA is re-accumulated through CLK1-dependent re-phosphorylation of SRSFs. The re-accumulation of the intron-retaining *CLK1* pre-mRNA is accelerated through the efficient re-phosphorylation within nSBs.^38,53,54^ As shown in Fig. 6b and 6c, the intron-retaining *CLK1* pre-mRNA was detected in normal conditions (lanes 1, 4 and 7) and converted to intron-excised mRNA upon thermal stress (lanes 2, 5 and 8) for the control and NCD- or QCD-treated cells. In contrast, re-accumulation of intron-retaining *CLK1* pre-mRNA was significantly suppressed only in the NCD-treated cells (lane 6, *cf*. lanes 3 and 9), suggesting that NCD specifically inhibits the nSB-dependent part of the auto-regulation mechanisms of *CLK1* pre-mRNA splicing. Consistently, NCD showed a limited effect on thermal stress-responsive alternative splicing of pre-mRNAs of *HSP105* and *TNRC6a* (Fig. 6b), which are unaffected by nSBs.

**Fig. 6.**
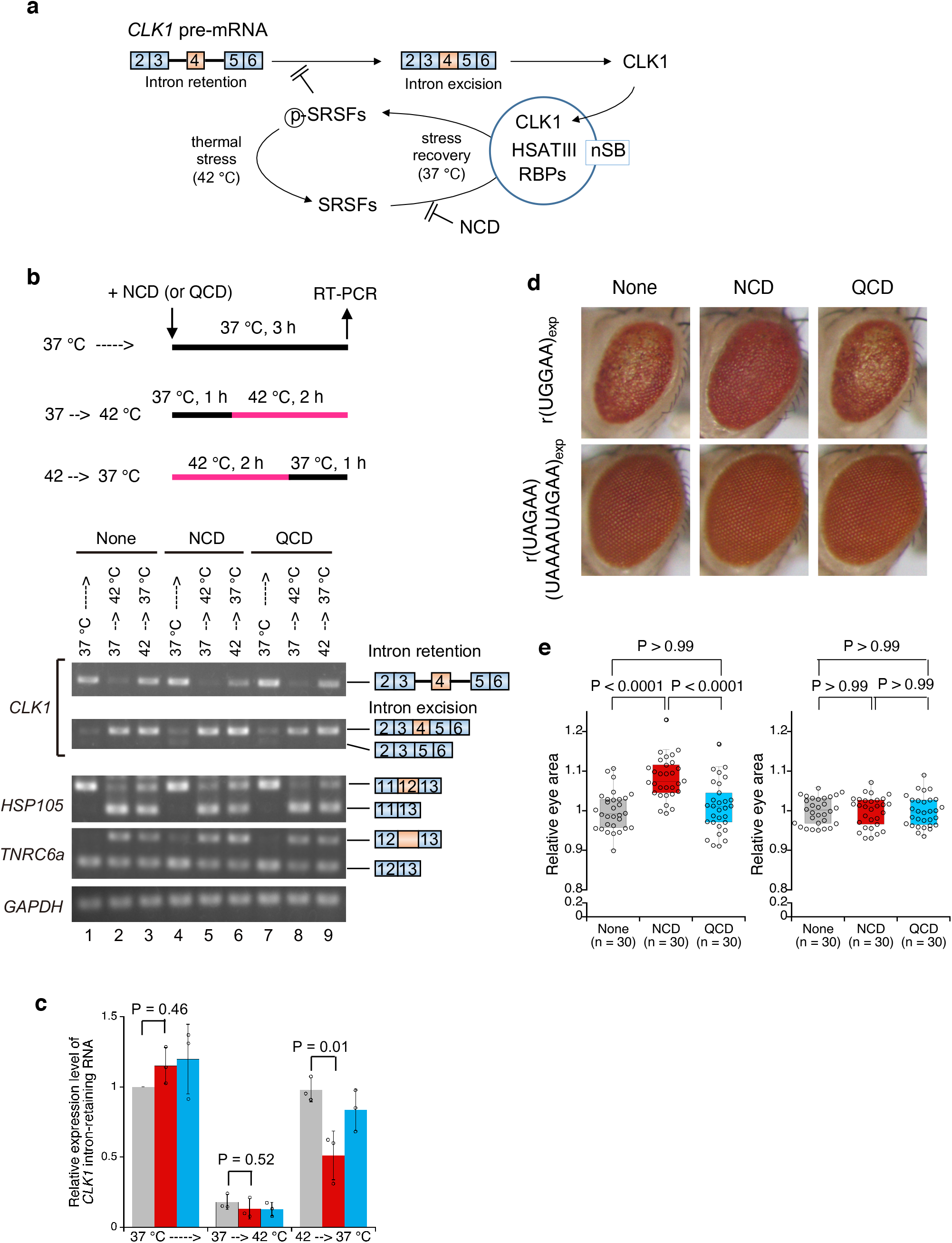
Biological activity of NCD in splicing and SCA31 disease models. **a**, Schematic illustration of nSB-dependent intron retention through re-phosphorylation of SRSFs. **b**, Semi-quantitative RT-PCR analysis of *CLK1* intron-retaining pre-mRNA during thermal stress recovery. The *GAPDH* mRNA was used as an internal control. **c**, Relative expression level of the *CLK1* intron-retaining pre-mRNA in the absence (gray) and presence of 2.5 μM NCD (red) and QCD (blue). Data are shown as the mean ± SD (n = 3); (Dunnett’s multiple comparison test). **d**, Light microscopic images of compound eyes from *Drosophila* expressing r(UGGAA)_exp_ (*GMR*-*Gal*4/+; *UAS*-(*UGGAA*)_exp_/+) (*upper, left*) and r(UAGAA)(UAAAAUAGAA)_exp_ (*GMR*-*Gal*4/+;; *UAS*-(*UAGAA*)(*UAAAAUAGAA*)_exp_/+) (*lower, left*) with the administration of NCD (middle) and QCD (right). Ligand concentration was 100 μM. **e**, Box plot of the eye area of *Drosophila* expressing r(UGGAA)_exp_ (left) and r(UAGAA)(UAAAAUAGAA)_exp_ (right). p-value (Kruskal-Wallis test, followed by Dunn’s multiple comparison test) was shown above.

### Effects of NCD on disease phenotype in *the Drosophila* model of SCA31

Having demonstrated that NCD disrupted r(UGGAA)_n_ RNA foci by interfering with RNA-protein interaction, we next investigated the biological activity of NCD in the *Drosophila* model of SCA31.^36^ This *Drosophila* model of SCA31 expresses pathogenic r(UGGAA)_80–100_ repeats (r(UGGAA)_exp_) in the compound eyes and showed degenerated eye morphology with depigmentation and decrease of the eye area (Fig. 6d; upper panel). *Drosophila* that expresses non-pathogenic repeats including r(UAGAA)_n_ and r(UAAAAUAGAA)_n_ was used as a control (Fig. 6d; lower panel, r(UAGAA)(UAAAAUAGAA)_exp_). When NCD (100 μM) was fed to larvae of the *Drosophila* model of SCA31, the degeneration of compound eyes in adults was suppressed (Fig. 6d; upper panel, middle image). The changes in the relative eye area between the NCD-treated and non-treated *Drosophila* expressing r(UGGAA)_exp_ were statistically significant (Fig. 6e; left). In marked contrast, QCD did not show any significant changes both in the morphology of compounds eye (Fig. 6d; upper panel, right image) and the eye area (Fig. 6e; left). For the control, *Drosophila* expressing r(UAGAA)(UAAAAUAGAA)_exp_ was affected neither by NCD nor QCD at all regarding morphology and eye area of compounds eye (Fig. 6d lower panel, and Fig. 6e right).

## Discussion

As the roles of functional non-coding RNAs (ncRNAs) in biological phenomenon and diseases became apparent, much needs for the small molecules binding to biologically relevant RNAs appeared. Synthetic RNA-binding molecules increase their importance in consideration of a growing number of evidence that various functional ncRNAs are involved in biological phenomenon and diseases. Among diverse approaches to discover RNA-binding molecules, our group has taken a repositioning approach of MBLs. MBLs consisted of two heterocycles having hydrogen bonding groups located at the edge of the heterocyclic ring, and the heterocycle can form the hydrogen-bonded pair with the specific nucleotide base. The hydrogen-bonded pairs would be further stabilized by stacking with the neighboring base pairs within the interior of the DNA π-stack.^26^ The molecule NCD we described here showed K_d(app)_ of 180 nM to the G-G mismatched base pair in dsDNA with the sequence context of 5’-CGG-3’/3’-GGC-5’ (Supplementary Fig. 13), but much decreased affinity was observed for the G-G mismatches (5’-CGG-3’/3’-GGC-5’ and 5’-GGC-3’/3’-CGG-5’) in dsRNA (data not shown). In contrast, K_d(app)_ for the NCD-binding to UGGAA/UGGAA pentad in dsRNA was 32 nM (Fig. 2d), which was 6-fold lower as compared with K_d(app)_ for the binding to 5’-CGG-3’/3’-GGC-5’ in dsDNA.

One of the common structural characteristics observed for the toxic RNAs is the formation of hairpin structures involving multiple mismatched base pairs.^49^ The expanded r(UGGAA)_n_ repeat RNA, causing SCA31 likely takes various secondary and tertiary structures, and these structures are under the dynamic equilibrium. We made use of our in-house MBL library containing heterocycles with different hydrogen bonding surface and identified NCD as the hit compound binding to the r(UGGAA)_n_ repeat RNA. The binding site of NCD in r(UGGAA)_n_ repeat RNA was firmly determined to be the UGGAA/UGGAA pentad in the secondary hairpin structure by NMR structural analysis, which also showed the substantial structural changes induced on the RNA. When U-A base pairs in the UGGAA/UGGAA pentad were replaced with C-G base pairs (i.e. CGGAG/CGGAG pentad), the binding of NCD to CGGAG/CGGAG pentad was comparable with UGGAA/UGGAA pentad (data not shown), suggesting that the U-A base pairs are not contributed to the NCD-binding. In addition, as NCD did not show the binding to G-G mismatch in 5’-GGC-3’/3’-CGG-5’-containing dsRNA (data not shown), the 5’-GGA-3’/3’-AGG-5’ internal loop structure is indispensable for the NCD-binding. In the NCD-SCA31 RNA complex, NCD bound to the RNA from the minor groove side. Each one of four guanines in the UGGAA/UGGAA pentad formed the hydrogen bonds to the *N*-acyl-2-amino-1,8-naphthyridine in NCD, and the produced guanine-naphthyridine hydrogen-bonded pairs were stacked inside the π-stack in dsRNA. Remarkably, all four guanine residues bound by naphthyridine formed a *syn* conformation regarding the glycosidic bond. Previously, we reported the NMR structure of the complex of CAG/CAG triad DNA with two molecules of naphthyridine-azaquinolone (NA), where the guanine-naphthyridine and adenine-azaquinolone hydrogen-bonded pairs were produced.^28^ In the case of the NA-CAG/CAG triad complex, NA bound to DNA from the major groove side and the glycosidic bond in guanosine and adenosine bound by NA adopt the *anti* conformation.

Our in vitro pulldown assay showed that NCD inhibited RNA-protein interaction between r(UGGAA)_n_ repeat RNAs and the RBPs, including TDP-43, HNRNPM, and SRSF9 that preferentially bind the r(UGGAA)_n_ repeat sequence. Previous studies reported that GAAUG was identified as a TDP-43-binding motif.^55^ Furthermore, it has been reported that HNRNPM and SRSF9 preferentially bound to purine-rich sequences.^55,56^ Our NMR structural analysis of the NCD-UGGAA/UGGAA complex indicated the NCD-binding to 5’-GGA-3’/3’-AGG-5’ internal loop through sequestration of four guanines. Notably, the recognition sequence of NCD in r(UGGAA)_n_ is overlapped with those of TDP-43, HNRNPM, and SRSF9, supporting the inhibitory effect of NCD on the specific interaction of r(UGGAA)_n_ with these RBPs. While a moderate decrease in pulldown efficiency was observed in FUS, it is likely due to multiple RNA-binding modes of FUS by sequence and shape recognition through two separate domains.^57^

We further investigated the mode of action of NCD to r(UGGAA)_n_ using the thermal stress-induced nSBs consisting of HSATIII lncRNAs as a model system, demonstrating that NCD impaired the nSB assembly without affecting HSATIII lncRNA levels. It has been shown that the RBP(s) harboring intrinsically disordered regions such as FUS and TDP-43 play a crucial role in the assembly of specific RNA foci.^58,59^ Therefore, the effect of NCD was attributed to impairment of RNA-protein interactions required for the nSB assembly that consequently leads to suppression of the nSB-dependent intron retention that we recently reported.^38^

Our studies on biological activity using the SCA31 models demonstrated reduction of the RNA foci in r(UGGAA)_n_-expressing cells and alleviation of the disease phenotype in the *Drosophila* model of SCA31. As far as we know, this is the first report of the small molecule that alleviates the SCA31 disease phenotype. It is remarkably noticeable that the difference in the effect of NCD and QCD on the SCA31 disease phenotype strongly correlates with their differences in the results of our *in vitro* binding, pulldown and cell-based assays. Although the elucidation of precise mechanisms alleviating the disease phenotype by NCD needs further studies, the disruption of r(UGGAA)_n_ RNA foci in the SCA31 cell model by NCD likely occurs in a similar mode of action to that of nSBs. Preceding studies demonstrated that accumulation of r(UGGAA)_n_ RNA foci clearly correlated with the severity of disease phenotype in *Drosophila* model of SCA31.^36^ In addition, the disruption of the RNA foci by binding of TDP-43 to r(UGGAA)_n_ alleviated the disease phenotype in *Drosophila* model of SCA31.^36^ The high affinity and specific binding of NCD to SCA31 repeat RNA may prevent pathogenic functions in r(UGGAA)_n_ such as RNA misfolding and RNA-protein interaction, eventually leading to the rescue of phenotype in *Drosophila* model of SCA31. We cannot exclude concerns regarding the off-target effect of NCD on other G-rich sequences existing in the human genome and transcripts because NCD was originally designed to bind to the G-rich sequences. However, the phenotypic change in *Drosophila* model of SCA31 by NCD is most likely due to the NCD-binding to r(UGGAA)_n_, because of high binding affinity of NCD to the UGGAA/UGGAA pentad, specific suppression of nSB-dependent splicing event by NCD, and the structure-activity relationship between NCD and QCD in the phenotypic change of *Drosophila* model of SCA31. Our demonstration of the mode of NCD-action to nSBs and the phenotype changes in the *Drosophila* model of SCA31 by NCD provides a potential therapeutic strategy of SCA31 for targeting r(UGGAA)_n_ RNAs by small molecules. The well-defined structures of NCD-RNA complex and bioactivity of NCD for the SCA31 disease model suggest that targeting mismatched base pairs in toxic repeat RNAs by small molecules could be a potential therapeutic strategy to treat repeat expansion diseases.

## Data availability

Solution structure of the complex of naphthyridine carbamate dimer and an RNA with UGGAA-UGGAA pentad have been deposited with Protein Data Bank (https://www.rcsb.org) under accession number 6IZP.

## References

1. Pearson, C. E., Edamura, K. N. & Cleary J. D. Repeat instability: Mechanisms of dynamic mutations. Nat. Rev. Genet. 6, 729–742 (2005).

2. Mirkin, S. M. Expandable DNA repeats and human disease. Nature 447, 932–940 (2007).

3. Orr, H. T. & Zoghbi, H. Y. Trinucleotide repeat disorders. Annu. Rev. Neurosci. 30, 575–621 (2007).

4. Liquori, C. L. et al. Myotonic dystrophy type 2 caused by a CCTG expansion in intron 1 of ZNF9. Science 293, 864–867 (2001).

5. DeJesus-Hernandez, M. et al. Expanded GGGGCC hexanucleotide repeat in noncoding region of C9ORF72 causes chromosome 9p-linked FTD and ALS. Neuron 72, 245–256 (2011).

6. Majounie, E. et al. Frequency of the C9orf72 hexanucleotide repeat expansion in patients with amyotrophic lateral sclerosis and frontotemporal dementia: a cross-sectional study. Lancet Neurol. 11, 323–330 (2012).

7. Matsuura, T. et al. Large expansion of the ATTCT pentanucleotide repeat in spinocerebellar ataxia type 10. Nat. Genet. 26, 191–194 (2000).

8. Kobayashi, H. et al. Expansion of intronic GGCCTG hexanucleotide repeat in NOP56 causes SCA36, a type of spinocerebellar ataxia accompanied by motor neuron involvement. Am. J. Hum. Genet. 89, 121–130 (2011).

9. Ishiura, H. et al. Expansion of intronic TTTCA and TTTTA repeats in benign adult familial myoclonic epilepsy. Nat. Genet. 50, 581–590 (2018).

10. Todd, P. K. & Paulson, H. L. RNA mediated neurodegeneration in repeat expansion disorders. Ann. Neurol. 67, 291–300 (2010).

11. Swinnen, B., Robberecht, W. & Bosch, L. V. D. RNA toxicity in non-coding repeat expansion disorders. EMBO J. e101112 (2019).

12. Jain, A. & Vale, R. D. RNA phase transitions in repeat expansion disorders. Nature 546, 243–247 (2017).

13. Sellier, C. et al. rbFOX1/MBNL1 competition for CCUG RNA repeats binding contributes to myotonic dystrophy type 1/type 2 differences. Nat. Commun. 9, 2009 (2018).

14. Zhang, Y.-J. et al. Heterochromatin anomalies and double-stranded RNA accumulation underlie C9orf72 poly(PR) toxicity. Science 336, eaav2606 (2019).

15. Zu, T. et al. Non-ATG–initiated translation directed by microsatellite expansions. Proc. Natl. Acad. Sci. USA 108, 260–265 (2011).

16. Mori, K. et al. The C9orf72 GGGGCC repeat is translated into aggregating dipeptide-repeat proteins in FTLD/ALS. Science 339, 1335–1338 (2013).

17. Krishnamurthy, M., Schirle, N. T. & Beal, P. A. Screening helix-threading peptides for RNA binding using a thiazole orange displacement assay. Bioorg. Med. Chem. 16, 8914–8921 (2008).

18. Zhang, J., Umemoto, S. & Nakatani, K. Fluorescent indicator displacement assay for ligand–RNA interactions. J. Am. Chem. Soc. 132, 3660–3661 (2010).

19. Fukuzumi, T., Murata, A., Aikawa, H., Harada, Y. & Nakatani, K. Exploratory study on the RNA-binding structural motifs by library screening targeting pre-miRNA-29a. Chem. Eur. J. 21, 16859–16867 (2015).

20. Patwardhan, N. N., Cai, Z., Newson, C. N. & Hargrove, A. E. Fluorescent peptide displacement as a general assay for screening small molecule libraries against RNA. Org. Biomol. Chem., 17,1778–1786 (2019).

21. Asare-Okai, P. N. & Chow, C. S. A modified fluorescent indicator displacement assay for RNA ligand discovery. Anal. Biochem. 408, 269–276 (2011).

22. Childs-Disney, J. L., Wu, M., Puschechnikov, A., Aminova, O. & Disney, M. D. A small molecule microarray platform to select RNA internal loop–ligand interaction. ACS Chem. Biol. 2, 745–754 (2007).

23. Sztuba-Solinska, J. et al. Identification of biologically active, HIV TAR RNA-binding small molecules using small molecule microarrays. J. Am. Chem. Soc. 136, 8402–8410 (2014).

24. Palacino, J. et al. SMN2 splice modulators enhance U1-pre-mRNA association and rescue SMA mice. Nat. Chem. Biol. 11, 511–517 (2015).

25. Howe, J. A. et al. Selective small-molecule inhibition of an RNA structural element. Nature 526, 672–677 (2015).

26. Nakatani, K. Recognition of Mismatched Base Pairs in DNA. Bull. Chem. Soc. Jpn. 82,1055–1069 (2009).

27. Nakatani, K., Sando, S. & Saito, I. Scanning of guanine-guanine mismatches in DNA by synthetic ligands using surface plasmon resonance assay. Nat. Biotechnol. 19, 51–55 (2001).

28. Nakatani, K. et al. Small-molecule ligand induces nucleotide flipping in (CAG)n trinucleotide repeats. Nat. Chem. Biol. 1, 39–43 (2005).

29. Peng, T. & Nakatani, K. Binding of naphthyridine carbamate dimer to the (CGG)n repeat results in the disruption of the G–C base pairing. Angew. Chem. Int. Ed. 44, 7280–7283 (2005).

30. Hagihara M., He H., Kimura M. & Nakatani K. A. Small Molecule Regulates Hairpin Structures in d(CGG) Trinucleotide Repeats. Bioorg. Med. Chem. Lett. 22, 2000–2003 (2012).

31. Li, J. et al. Naphthyridine-benzoazaquinolone: evaluation of a tricyclic system for the binding to (CAG)n repeat DNA and RNA. Chem. Asian J. 11, 1971–1981 (2016).

32. Matsumoto, J., Li, J., Dohno, C. & Nakatani, K. Synthesis of 1H-pyrrolo[3,2-h]quinoline-8-amine derivatives that target CTG trinucleotide repeats. Bioorg. Med. Chem. Lett. 26, 3761–3764 (2016).

33. Ishikawa, K. et al. Pentanucleotide repeats at the spinocerebellar ataxia type 31 (SCA31) locus in Caucasians. Neurology. 77, 1853–1855 (2011).

34. Sato, N. et al. Spinocerebellar ataxia type 31 is associated with ‘‘inserted’’ penta-nucleotide repeats containing (TGGAA)n. Am. J. Hum. Genet. 85, 544–557 (2009).

35. Niimi, Y. et al. Abnormal RNA structures (RNA foci) containing a penta-nucleotide repeat (UGGAA)n in the Purkinje cell nucleus is associated with spinocerebellar ataxia type 31 pathogenesis. Neuropathology 33, 600–611 (2013).

36. Ishiguro, T. et al. Regulatory role of RNA chaperone TDP-43 for RNA misfolding and repeat-associated translation in SCA31. Neuron 94, 108–124 (2017).

37. Denegri, M. et al. Human Chromosomes 9, 12, and 15 contain the nucleation sites of stress-induced nuclear bodies. Mol. Biol. Cell 13, 2069–2079 (2002).

38. Ninomiya, K et al. LncRNA-dependent nuclear stress bodies promote intron retention through SR protein phosphorylation. EMBO J. e102729 (2019).

39. Warf, M. B., Nakamori, M., Matthys, C. M., Thornton, C. A. & Berglund, J. A. Pentamidine reverses the splicing defects associated with myotonic dystrophy. Proc. Natl. Acad. Sci. USA 106, 18551–18556 (2009).

40. Nguyen, L. et al. Rationally designed small molecules that target both the DNA and RNA causing myotonic dystrophy type 1. J. Am. Chem. Soc. 137, 14180–14189 (2015).

41. Rzuczek, S. G. et al. Precise small-molecule recognition of a toxic CUG RNA repeat expansion. Nat. Chem. Biol. 13, 188–193 (2017).

42. Rzuczek, S. G., Park, H. & Disney, M. D. A toxic RNA catalyzes the in cellulo synthesis of its own inhibitor. Angew. Chem. Int. Ed. 53, 10956–10959 (2014).

43. Nguyen, L., Lee J. Y. & Zimmerman, S. C. Small molecules that target the toxic RNA in myotonic dystrophy type 2. ChemMedChem. 9, 2455–2462 (2014).

44. Yang, W.-Y., Wilson, H. D., Velagapudi, S. P. & Disney, M. D. Inhibition of non-ATG translational events in cells via covalent small molecules targeting RNA. J. Am. Chem. Soc. 137, 5336–5345 (2015).

45. Yang, W.-Y. et al. Small molecule recognition and tools to study modulation of r(CGG^exp^ in fragile X-associated tremor ataxia syndrome. ACS Chem. Biol. 11, 2456–2465 (2016).

46. Su, Z. et al. Discovery of a biomarker and lead small molecules to target r(GGGGCC)-associated defects in c9FTD/ALD. Neuron 83, 1043–1050 (2014).

47. Simone, R. et al. G-quadruplex-binding small molecules ameliorate C9orf72 FTD/ALS pathology in vitro and in vivo. EMBO Mol. Med. 10, 22–31 (2018).

48. Yang, W.-Y., Gao, R., Southern, M., Sarkar, P. S. & Disney, M. D. Design of a bioactive small molecule that targets r(AUUCU) repeats in spinocerebellar ataxia 10. Nat. Commun. 7, 11647 (2016).

49. Ciesiolka, A., Jazurek, M., Drazkowska, K. & Krzyzosiak, W. J. Structural characteristics of simple RNA repeats associated with disease and their deleterious protein interactions. Front. Cell. Neurosci. 11, 97 (2017).

50. Davis, I. W. et al. MolProbity: All-atom contacts and structure validation for proteins and nucleic acids. Nucleic Acids Res. 35, 375–383 (2007).

51. Aly, M. K., Ninomiya, K., Adachi, S., Natsume, T. & Hirose, T. Two distinct nuclear stress bodies containing different sets of RNA-binding proteins are formed with HSATIII architectural noncoding RNAs upon thermal stress exposure. Biochem. Biophys. Res. Commun. 516, 419–423 (2019).

52. Jolly, C et al. Stress-induced transcription of satellite III repeats. J. Cell Biol. 164, 25–33 (2004).

53. Duncan, P. I. et al. Alternative splicing of STY, a nuclear dual specificity kinase. J. Biol. Chem. 270, 21524–21531 (1995).

54. Ninomiya, K., Kataoka, N. & Hagiwara, M. Stress-responsive maturation of Clk1/4 pre-mRNAs promotes phosphorylation of SR splicing factor. J. Cell Biol. 195, 27–40 (2011).

55. Ray, D. et al. A compendium of RNA-binding motifs for decoding gene regulation. Nature 499, 172–177 (2013).

56. Cho, S. et al. hnRNP M facilitates exon 7 inclusion of SMN2 pre-mRNA in spinal muscular atrophy by targeting an enhancer on exon 7. Biochim. Biophys. Acta 1839, 306–315 (2014).

57. Loughhlin, F. E. et al. The solution structure of FUS bound to RNA reveals a bipartite mode of RNA recognition with both sequence and shape specificity. Mol. Cell 73, 490–504 (2019).

58. Henning, S. et al. Prion-like domains in RNA binding proteins are essential for building subnuclear paraspeckles. J. Cell Biol. 210, 529–539 (2015).

59. Mittag, T. & Parker, R. Multiple modes of protein-protein interactions promote RNP granule assembly. J. Mol. Biol. 430, 4636–4649 (2018).

