## Supplementary Information for "Small molecule targeting r(UGGAA)_n_ disrupts RNA foci and alleviates disease phenotype in *Drosophila* model"

### Contents

| Methods | | P 3–8 |
|---|---|---|
| **Supplementary Figure 1** | Chemical structures of 20 compounds (LC-1–20) in in-house chemical library. | P 9 |
| **Supplementary Figure 2** | SPR analysis of the binding of LC-1–20 to r(UGGAA)$_9$-, r(UAGAA)$_9$- and r(UAAAA)$_9$-immobilized surfaces. | P 10 |
| **Supplementary Figure 3** | EMSA to confirm interactions of r(UGGAA)$_9$ with molecules. | P 11 |
| **Supplementary Figure 4** | EMSA to confirm interactions of r(UGGAA)$_9$ with molecules. | P 12 |
| **Supplementary Figure 5** | Thermal melting curves of r(UAGAA)$_9$ and r(UAAAA)$_9$ in the absence and presence of NCD. | P 13 |
| **Supplementary Figure 6** | Thermal melting curves of RRA/RRA internal loop-containing RNA duplexes in the absence and presence of NCD or QCD. | P 14 |
| **Supplementary Figure 7** | Thermal melting curves and CD spectra of UGGAA-UGGAA pentad-containing hairpin RNA. | P 15 |
| **Supplementary Figure 8** | ESI-TOF-MS analysis of UGGAA-UGGAA pentad-containing hairpin RNA with NCD. | P 16 |
| **Supplementary Figure 9** | Examples of intermolecular NOEs of SCA31RNA and NCD. | P 17 |
| **Supplementary Figure 10** | EMSA to confirm the interaction of r(UGGAA)$_9$ with RNA FISH probe in the absence and presence of NCD and WST-1 assay and qPCR of r(UGGAA)$_{76}$ in the absence and presence of ligands. | P 18 |
| **Supplementary Figure 11** | RNA FISH and IF images of HeLa cells expressing r(UGGAA)$_{76}$ in the absence and the presence of NCD or QCD stained by r(UGGAA)$_{76}$-FISH and IF using anti-TDP-43 antibodies. | P 19 |
| **Supplementary Figure 12** | RNA FISH and IF images of HeLa cells after thermal stress exposure in the absence and the presence of NCD or QCD stained by HSATIII-FISH and IF using anti-SRSF9 antibodies. | P 20 |
| **Supplementary Figure 13** | ITC measurements for the binding of NCD to CGG/CGG-containing hairpin DNA. | P 21 |
| **Supplementary Table 1** | NMR constraints and structure statistics. | P 22 |
| **Supplementary Table 2** | All-atom structure validation by MolProbity | P 23–24 |
| **Supplementary Table 3** | Primer sequences used in quantitative and semi-quantitative RT-PCR | P 25 |

### Methods

#### SPR analyses

5’-biotin-TEG r(UGGAA)$_9$, r(UAGAA)$_9$ and r(UAAAA)$_9$ repeat RNAs were immobilized on the SA sensor chip (BIAcore) that coated the surface with streptavidin. The surface of sensor chip SA was washed with 50 mM NaOH and 1 M NaCl at three times for 60 s with the flow rate of 30 mL min$^{-1}$. 5’-biotin-TEG repeat RNAs was immobilized to the surface under the following conditions: 200 nM repeat RNA in 10 mM HEPES (pH 7.4), 500 mM NaCl. Amount of r(UGGAA)$_9$, r(UAGAA)$_9$ and r(UAAAA)$_9$ immobilized on the chip surface was 586, 658, 529 response units (RU), respectively. SPR analysis for the binding of in-house chemical library to the repeat RNA-immobilized surfaces was performed using a BIAcore T200 SPR system (GE Healthcare) under the following condition: 500 nM compounds in HBS-EP+ buffer (GE Healthcare) containing 10 mM HEPES (pH7.4), 150 mM NaCl, 3 mM EDTA, 0.05% v/v Surfactant P20.

#### EMSA experiments

Repeat RNA (200 nM) without and with ligand (2 μM) in 10 mM sodium cacodylate buffer (pH 7.0) containing 100 mM NaCl was incubated at room temperature for 15 min. The mixtures were mixed with 10 × loading buffer (TAKARA) and were subjected to electrophoresis through a native polyacrylamide gel with 1× Tris/Borate/EDTA buffer at room temperature. After electrophoresis, the gels were stained for 10 min with SYBR Gold (Thermo). Images of the gels were collected using a Safe Imager 2.0 Blue Light Transilluminator (Thermo).

#### Melting temperature measurements

Thermal denaturation profiles were recorded on a UV-2700 spectrophotometer (Shimadzu) equipped with the TMSPC-8 temperature controller. The absorbance of RNA (4 μM for RNA duplexes and 2 μM for repeat RNAs) without and with ligand (20 μM) in 10 mM sodium cacodylate buffer (pH 7.0) containing 100 mM NaCl was monitored at 260 nm from 2 to 100 °C (1 °C min$^{-1}$). $T_m$ was calculated using the median method.

**CSI-TOF-MS measurements**

Samples were prepared by mixing UGGAA/UGGAA pentad-containing hairpin-RNA (10 μM) and NCD (5–40 μM) in 50% methanol in water containing 100 mM ammonium acetate. Mass spectra were obtained with JEOL JMS-T100LP AccuTOF LC-plus 4G mass spectrometer in negative mode. Spray temperature was fixed at –10 °C with a sample flow rate of 20 μL $min^{-1}$.

**CD measurements**

CD experiments were carried out on a J-725 CD spectrometer (JASCO) using a 10 mm path length cell. CD spectra of RNA duplex (4 μM) in the absence and presence of ligand (20 μM) were measured in 10 mM sodium cacodylate buffer (pH 7.0) containing 100 mM NaCl.

**ITC measurements**

A solution of UGGAA/UGGAA pentad-containing hairpin-RNA (2.5 μM) or CGG/CGG-containing hairpin DNA (4 μM) was titrated with NCD solution (50 μM for RNA or 80 μM for DNA) at 25 °C in 10 mM sodium cacodylate buffer (pH 7.0) containing 100 mM NaCl on a MicroCal $iTC_{200}$ calorimeter. Thermodynamic parameters were calculated from the binding curve using Microcal origin 7.0 with a binding model involving a single set of identical sites.

**NMR structural analysis**

RNA sample for NMR measurements was purchased from (GeneDesign). RNA sample was dissolved in 20 mM sodium phosphate buffer (pH 6.9) with 100 mM NaCl and 5% $D_2O$. The sample concentration was 0.075 mM. NMR spectra were measured at 288 K with Avance600 spectrometer (Bruker BioSpin). After measurement of one-dimensional as well as two-dimensional spectra including HOHAHA and NOESY. For water signal suppression, the jump-and-return pulse[1] was used for the 1D imino proton spectra, and the 3-9-19 pulse[2] were used for other measurements. After the NMR measurement in free form, the SCA31 hpRNA was titrated by NCD with molar ratio of 1:1.5, 1:2, 1:2.5, 1:2.75, 1:3, and 1:3.5. Then, the excess amount of NCD was removed by ultrafiltration by Vivaspin (Amicon). The solvent was replaced by 100% $D_2O$, and NMR spectra were measured for structural determination.

NMR data were processed with TopSpin (Bruker Biospon) and analyzed with Sparky.[3] Most of the signals for base protons and H1' protons were assigned for RNA, and all CH protons of NCD were assigned. Structure of the RNA-NCD complex was calculated with CNS_SOLVE.[4] 192 distance restraints including 36 hydrogen bonding restraints, 157 dihedral restraints, and 14 planarity restraints were used (Supplementary Table 1). The geometry and atomic charge for NCD in an extended conformation was prepared with UCSF Chiemra[5] and ANTECHAMBER.[6]

Forty-four structures without any violation for restraints were obtained by 100 calculations. Ten structures with the lowest energy were chosen, and the averaged-minimized structure was obtained. R.m.s.d. for the ten structures for heavy atoms was 1.966 ± 0.775 and 1.069 ± 0.388 Å for all and the core region, U5-G7, A9, U21-G23, A25, and NCD, respectively.

**Plasmid construction**

Preparation of plasmid containing $(TGGAA)_{76}$ was delegated to Thermo Fisher Scientific. The $(TGGAA)_{76}$-containing plasmid was digested with *Eco*RI and *Xba*I, and inserted into *Eco*RI–*Xba*I site of pCMVTnT to give pCMVTnT-$(TGGAA)_{76}$ plasmid.

**Cell culture**

HeLa cells (originally obtained from RIKEN BRC) were cultured at 37 °C in 5% $CO_2$ in Dulbecco's modified eagle's medium (Sigma) containing 10% fetal bovine serum (MP Biomedicals), penicillin and streptomycin (Thermo). For thermal stress induction, the cells were incubated at 42 °C in an incubator with 5% $CO_2$.

***In vitro* pulldown assay**

In vitro pulldown assay was performed as previously reported.[7] In brief, $(GGAAT)_{20}$ dsDNA was introduced into pCR-Blunt II-TOPO vector (Thermo Fisher). Then, UGGAA and the antisense repeat RNAs were synthesized by in vitro transcription using T7 and SP6 RNA polymerases, respectively with Biotin RNA labeling mix (sigma) and purified using Centri-Sep Spin Columns (Thermo Fisher). Biotinylated $r(UGGAA)_{20}$ or $r(UUCCA)_{20}$ repeat RNA (1 μg) was bound to Tamavidin2-REV magnetic beads, mixed with pre-absorbed HeLa nuclear extract diluted in PBS containing 0.1% Triton X-100, 1× protease inhibitor, 1 mM PMSF and RNase inhibitor without and with NCD or QCD (2 μM), and rotated overnight at 4 °C. After washing five times with cold PBS containing 0.1% Triton X-100, the co-precipitated proteins were eluted in SDS sample buffer for 5 min at 95 °C.

**Western blotting**

Protein extracts in 1× SDS sample buffer were boiled for 5 min, separated by SDS-PAGE, and transferred to a PVDF membrane (Millipore) by electroblotting. After incubation with primary and HRP-conjugated secondary antibodies, the signals on the membranes were developed by a chemiluminescence reaction using the ImmunoStar Kit (Wako Chemicals), detected with a ChemiDoc imaging system (BioRad), and analyzed using ImageJ software (NIH). Antibodies are as follows. Anti-HNRNPM (LifeSpan Biosciences, LS-B2427), anti-SFPQ (MBL, RN014MW), anti-SRSF9 (MBL, RN081PW), anti-ALYREF (Santa cruz, sc323-11), anti-TDP-43 (Proteintech, 10782-2-AP), anti-FUS (Santa cruz, sc477-11), anti-Mouse IgG (HRP-Linked) (GE Healthcare, NA931-1ML), anti-Rabbit IgG (HRP-Linked) (GE Healthcare, NA934-1ML).

**RNA fluorescence *in situ* hybridization (FISH) and immunofluorescence (IF)**

A total of $1 \times 10^5$ cells were seeded into 24-well plates containing coverslips. After a 24 h culture, the cell culture medium containing compound was added to the cells. Plasmids (500 ng) expressing $r(UGGAA)_{76}$ were transfected to the cells using 2 μl of FuGENE HD Transfection Reagent (Promega). After 24 h of transfection, the cells were washed with phosphate-buffered saline (PBS) and fixed with 4% paraformaldehyde for 30 min at 4 °C. The fixed cells were washed with PBS and permeabilized with PBS containing 2% acetone pre-chilled at –20 °C for 5 min at 4 °C. Subsequently, the cells were washed with PBS and stored in 70% EtOH at –20 °C overnight. After washing with PBS and rehydration with 30% formamide in 2 × saline sodium citrate (SSC) for 10 min at room temperature, the cells were pre-hybridized in hybridization buffer (30% formamide, 2 × SSC, 66 μg/ml yeast tRNA, 0.02% BSA, 10% dextran sulfate, 2 mM Ribonucleoside-Vanadyl Complex) for 30 min at 37 °C and hybridized in hybridization buffer containing 1 nM Alexa647-labeled $(TTCCA)_5$ DNA/LNA probe (sequence detail was shown in Supplementary Fig. 12) for 2 h at 37 °C. The washing of the coverslips was performed three times in 2 × SSC containing 50% formamide, two times in 1 × SSC and two times in 0.1 × SSC for 20 min at 55 °C. The cells were mounted onto microscope slides with

SlowFade Diamond containing DAPI (Thermo). Fluorescence images of cells were taken by BZ-9000 Fluorescence Microscope (KEYENCE). FISH and IF staining of nSBs and nuclear speckles were performed as previously reported.[8] Antibodies used are as follows (also see western blotting section). Anti-SAFB (Abcam, ab8060), anti-SRSF2 (Sigma-Aldrich, S4045), anti-Digitonin (Abcam, ab420 and ab76907), anti-TDP-43 (Proteintech, 10782-2-AP), anti-Mouse IgG (H+L) (Alexa Fluor 488) (Thermo Fisher Scientific, A11029), anti-Rabbit IgG (H+L) (Alexa Fluor488) (Thermo Fisher Scientific, A11034), anti-Rabbit IgG (H+L) (Alexa Fluor568) (Thermo Fisher Scientific, A11036), anti-Goat IgG H&L (Alexa Flour 488) (Abcam, ab150129), anti-Mouse IgG H&L (Alexa Fluor 568) (Abcam, ab175472), . The image analysis was performed using ImageJ (https://imagej.nih.gov/ij/) software. Areas of r(UGGAA)$_{76}$ RNA foci, nSBs, nuclear speckles, and nuclei were defined and measured by binarized images of r(UGGAA)$_{76}$ RNA, HSATIII, SRSF2, and DAPI, respectively. These data of r(UGGAA)$_{76}$ RNA foci, nSBs, and nuclear speckles area in nucleus were shown as boxplot containing the minimum, the maximum, the median, and the first and third quartiles.

**Reverse transcription-quantitative PCR (RT-qPCR)**

RT-qPCR and semi-quantitative RT-PCR were performed as previously reported.[8] In brief, total RNAs were prepared using TRI Reagent (Molecular Research Center, Inc.), according to the manufacturer's manual. The RNAs were treated with RQ1 RNase-free DNase (Promega) according to the manufacturer's manual. First-strand cDNA was synthesized using High-Capacity cDNA Reverse Transcription Kits (Thermo Fisher Scientific). For RT-qPCR, the cDNAs were amplified using KAPA SYBR FAST qPCR Master Mix (KAPA Biosystems) and monitored using the LightCycler 480 System (Roche). For semi-quantitative RT-PCR**,** cDNAs were amplified by PCR to unsaturated levels, separated by electrophoresis, and stained with ethidium bromide. Images were obtained with a ChemiDoc system (BioRad) and analyzed with ImageJ software (NIH). Primers for PCR are shown in Supplementary Table 3.

**Cell viability assay**

Effect of small molecule treatment on cell viability was determined by WST-8 assay (CCK-8 kit, Dojindo). HeLa cells were seeded at $2 \times 10^4$ cells/well in 96-well plates and cultured for 24 h. Small molecules were added to final concentrations of 5 μM to the medium and the cells were cultured further for 24 h. CCK-8 solution (5 μl) was added to each well and the plates were further incubated for several hours at 37°C before the measurement of the absorbance at 450 nm by a plate reader (EL808, BioTek).

**Fly experiment**

Instant *Drosophila* Medium Blue containing dry yeast was mixed with ultrapure water or 100 μM compound. Parent flies (*GMR*-*Gal*4/+; and *UAS*-(*UGGAA*)$_{exp}$/+ for *Drosophila* expressing r(UGGAA)$_{exp}$ or *GMR*-*Gal*4/+;; and *UAS*-(*UAGAA*)(*UAAAAUAGAA*)$_{exp}$/+ for *Drosophila* expressing r(UAGAA)(UAAAAUAGAA)$_{exp}$) were crossed on the food without and with 100 μM compound, and the offspring were generated on the same food at 25 °C. The eye morphology of the 1-2-day-old flies was analyzed using the stereoscopic microscope model SZX10 (Olympus).

**References in Methods**

1. Plateau, P. & Gueron, M. Exchangeable proton NMR without base-line distortion, using new strong-pulse sequences. *J. Am. Chem. Soc.* **104**, 7310–7311 (1982).
2. Piotto, M., Saudek, V. & Sklenár, V., Gradient-tailored excitation for single-quantum NMR spectroscopy of aqueous solutions. *J. Biomol. NMR* **2**, 661–665 (1992).
3. Goddard, T.D. & Kneller, D.G. SPARKY 3, University of California, San Francisco
4. Brünger, A.T. *et al.* Crystallography and NMR System (CNS): a new software system for macromolecular structure determination. *Acta Cryst.* **D54**, 905–921 (1998).
5. Pettersen, E.F. *et al.* UCSF Chimera - a visualization system for exploratory research and analysis. *J. Comput. Chem.* **25**, 1605-1612 (2004).
6. Wang, J., Wang, W., Kollman, P. A. & Case, D. A., Automatic atom type and bond type perception in molecular mechanical calculations, *J. Mol. Graph. Model.* **25**, 247–260 (2006).
7. Yamazaki, T. *et al.* Functional domains of NEAT1 architectural lncRNA induce paraspeckle assembly through phase separation. *Mol. Cell* **70**, 1038–1053 (2018).
8. Ninomiya, K *et al.* LncRNA-dependent nuclear stress bodies promote intron retention through SR protein phosphorylation. *EMBO J.* e102729 (2019).

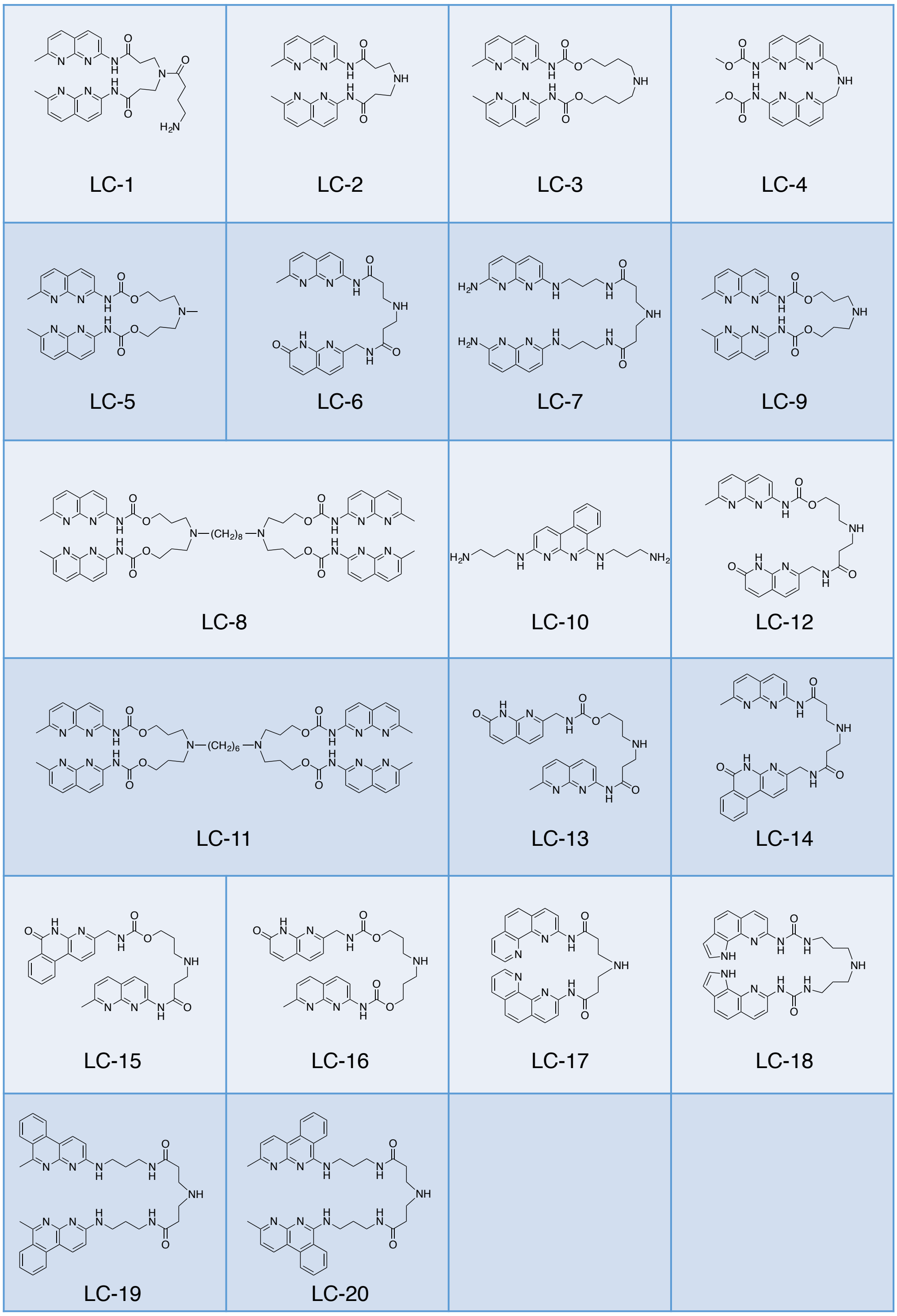

**Supplementary Figure 1**

Chemical structures of 20 compounds (LC-1–20) in in-house chemical library.

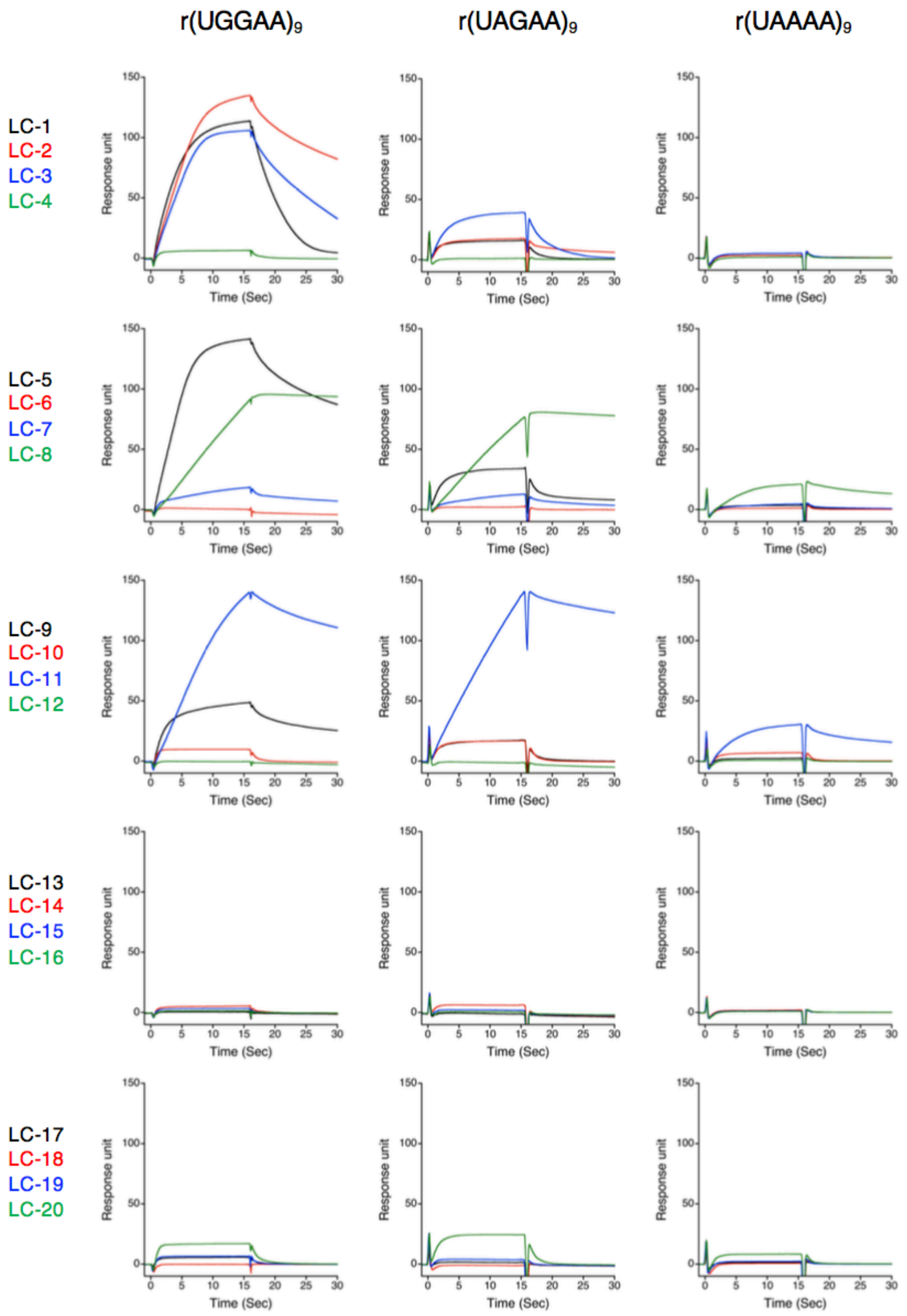

**Supplementary Figure 2**

SPR analysis of the binding of LC-1–20 to r(UGGAA)$_9$-, r(UAGAA)$_9$- and r(UAAAA)$_9$-immobilized surfaces. Compound concentration was 500 nM. The amount of 5’-Biotin-labelled r(UAGAA)$_9$ and r(UAAAA)$_9$ immobilized on the SA sensor chip was 658 and 528 RU, respectively.

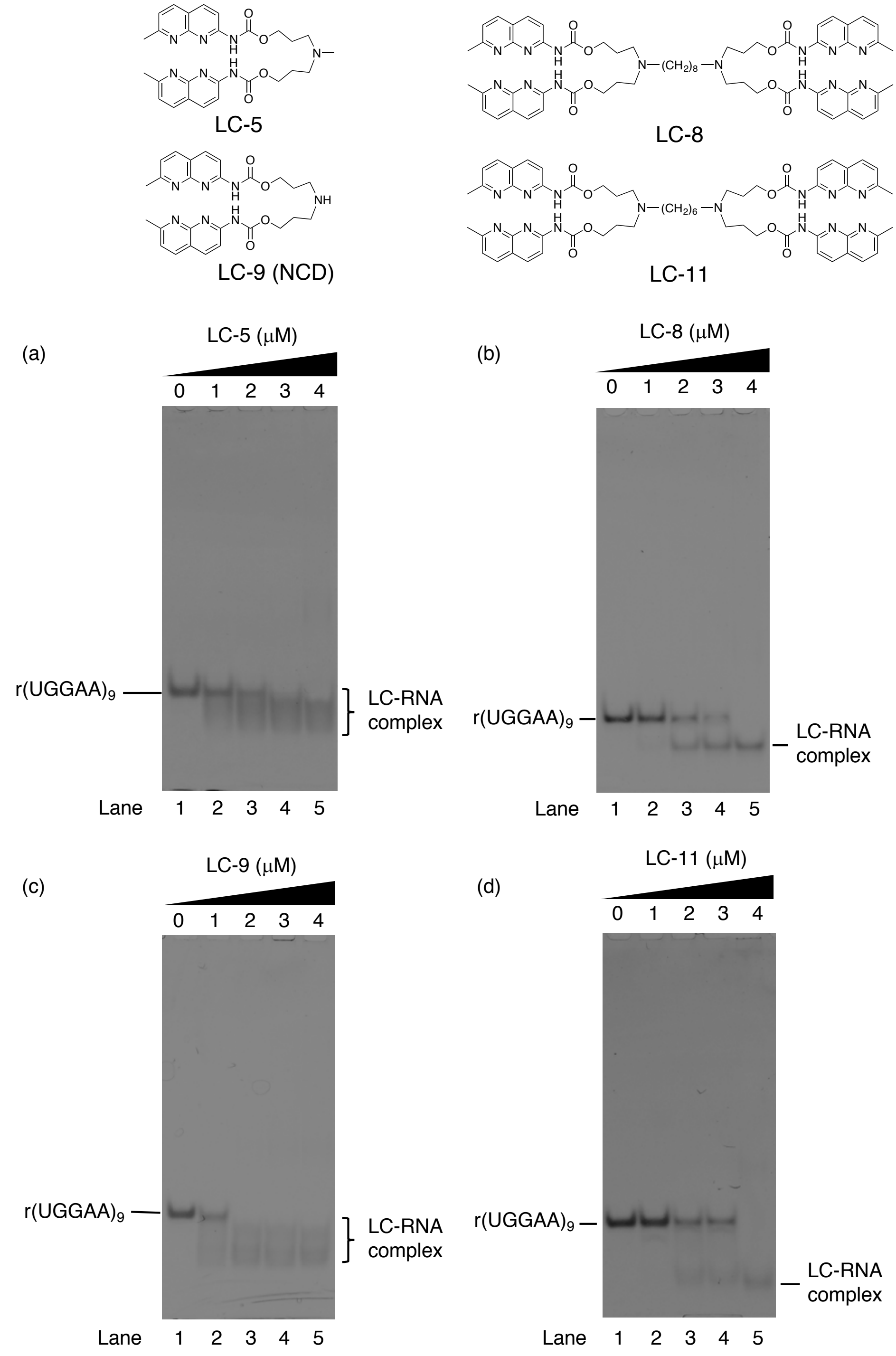

**Supplementary Figure 3**

EMSA to confirm interactions of r(UGGAA)$_9$ with (a) LC-5, (b) LC-8, (c) LC-9, and (d) LC-11. RNA concentration: 200 nM. Compound concentration was 0, 1, 2, 3 and 4 μM.

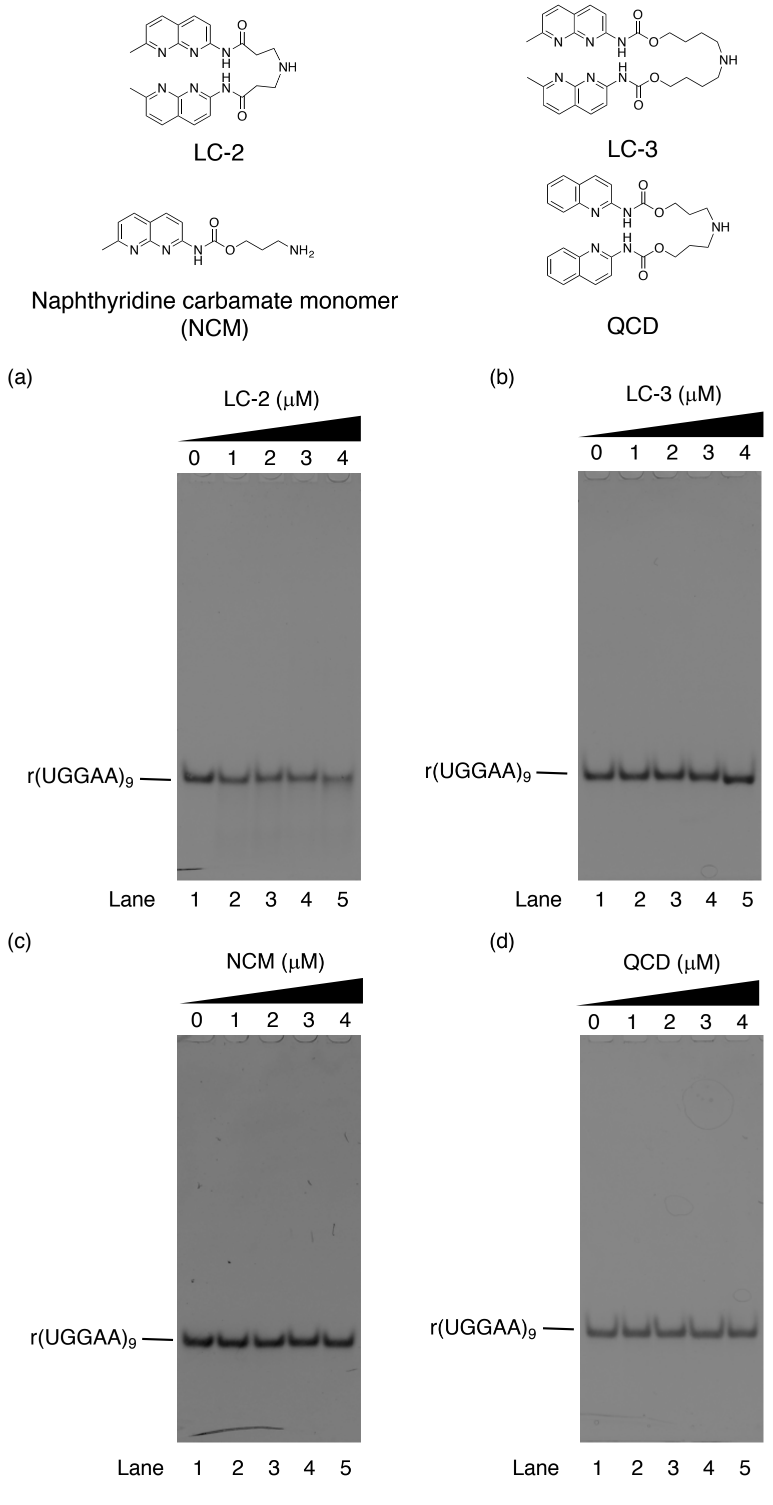

**Supplementary Figure 4**

EMSA to confirm interactions of r(UGGAA)$_9$ with (a) LC-2, (b) LC-3, (c) NCM, and (d) QCD. RNA concentration: 200 nM. Compound concentration was 0, 1, 2, 3 and 4 μM.

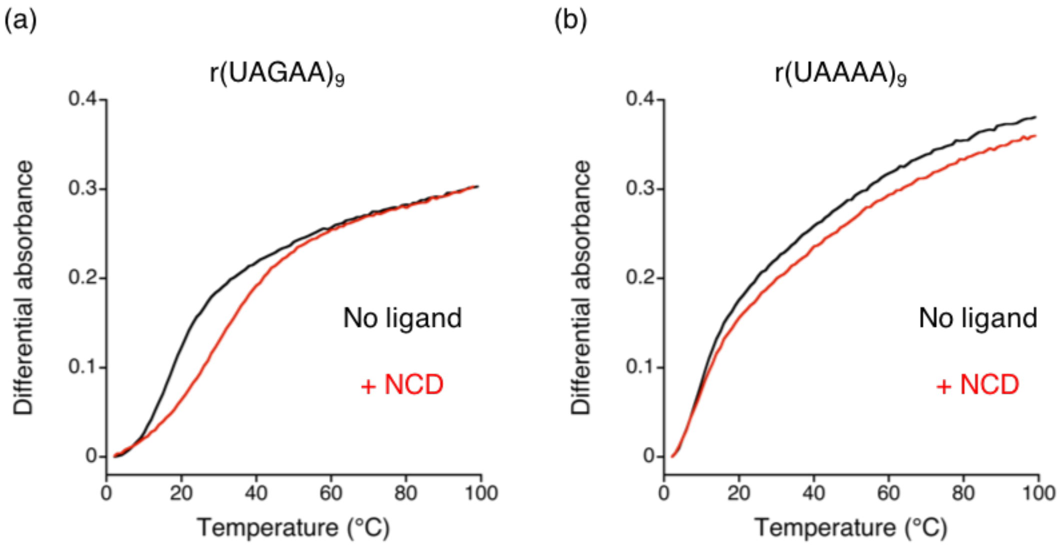

(c)

| | $T_m$ (°C) | |
|---|---|---|
| | No ligand | + NCD |
| r(UGGAA)$_9$ | N.D. | 73.7 (1.1) |
| r(UAGAA)$_9$ | N.D. | 31.1 (1.0) |
| r(UAAAA)$_9$ | N.D. | N.D. |

**Supplementary Figure 5**

Thermal melting curves of (a) r(UAGAA)$_9$ and (b) r(UAAAA)$_9$ (2 μM) in the absence (black) and presence of NCD (red) in 10 mM sodium cacodylate (pH 7.0) containing 100 mM NaCl. Ligand concentration was 20 μM. (c) Table of $T_m$ values of r(UGGAA)$_9$, r(UAGAA)$_9$, and r(UAAAA)$_9$ in the absence and presence of NCD (20 μM)

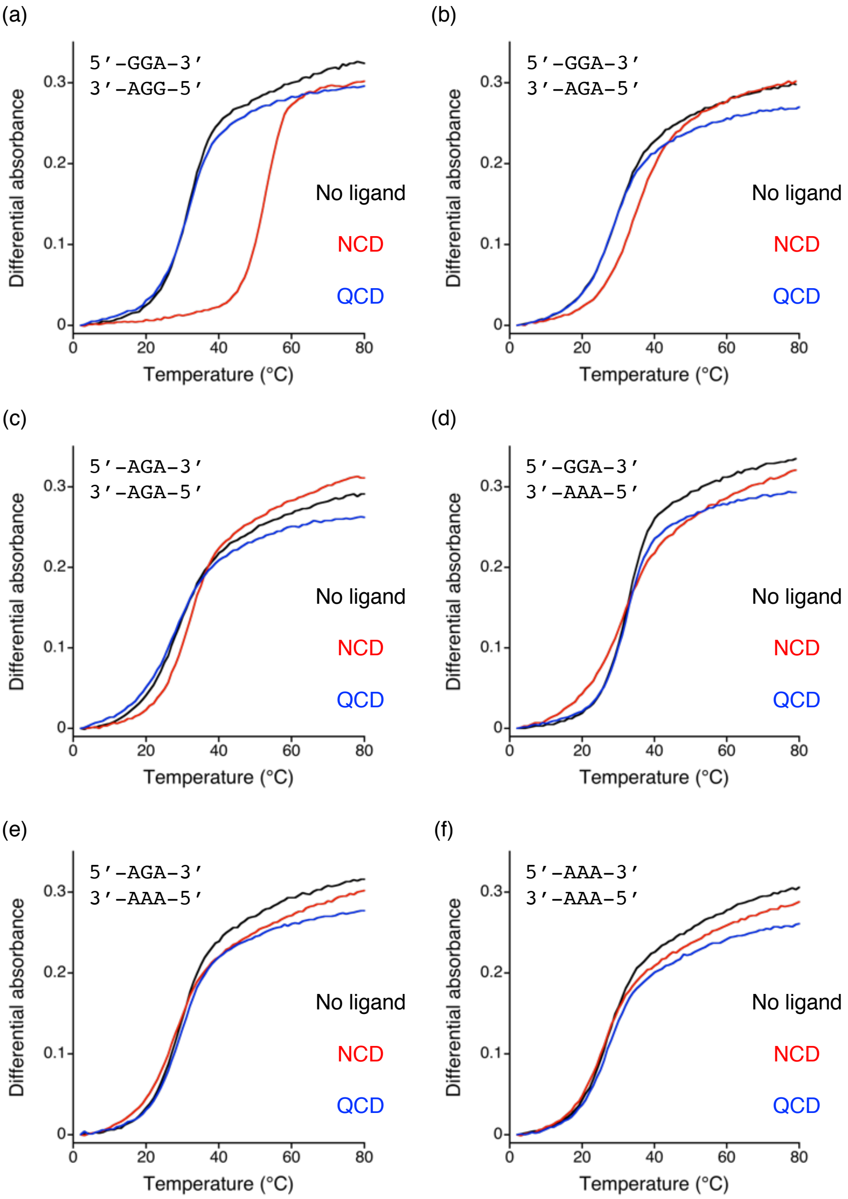

**Supplementary Figure 6**

Thermal melting curves of RRA/RRA internal loop-containing RNA duplexes (4 μM), r(GUACURRAACAUG)/r(CAUGURRAAGUAC) in the absence (black) and presence of NCD (red) or QCD (blue) in 10 mM sodium cacodylate (pH 7.0) containing 100 mM NaCl. (a) GGA/GGA, (b) GGA/AGA, (c) AGA/AGA, (d) GGA/AAA, (e) AGA/AAA, (f) AAA/AAA internal loop-containing RNA duplexes. Ligand concentration was 20 μM.

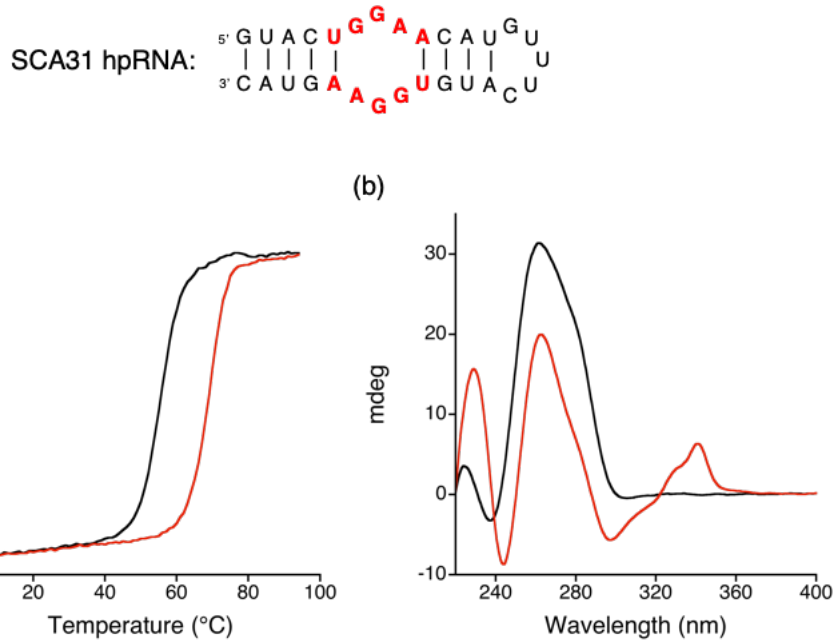

**Supplementary Figure 7**

(a) Thermal melting curves and (b) CD spectra of SCA31 hpRNA (4 μM), 5'-r(GUAC UGGAA CAUGUUUCAUG UGGAA GUAC)-3' without (black) and with (red) NCD (20 μM) in 10 mM sodium cacodylate (pH 7.0) containing 100 mM NaCl.

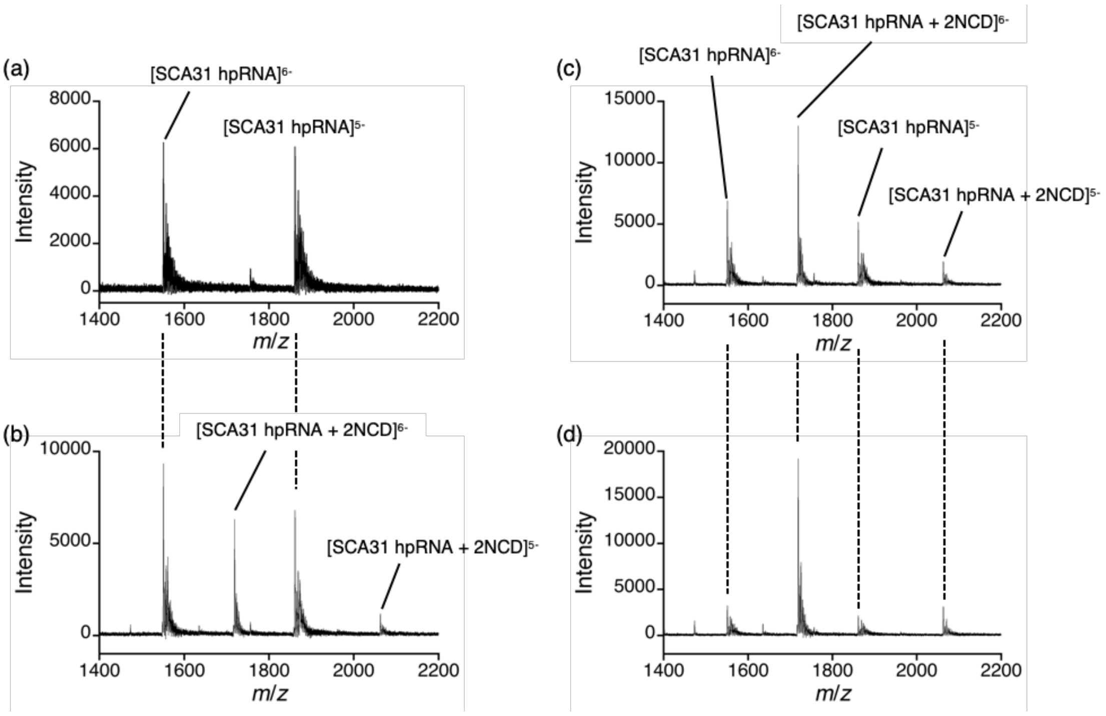

**Supplementary Figure 8**

ESI-TOF-MS analysis of SCA31 hpRNA (10 μM), 5'-r(GUAC UGGAA CAUGUUUCAUG UGGAA GUAC)-3' with NCD. Ligand concentration was (a) 0, (b) 5, (c) 10 and (d) 20 μM.

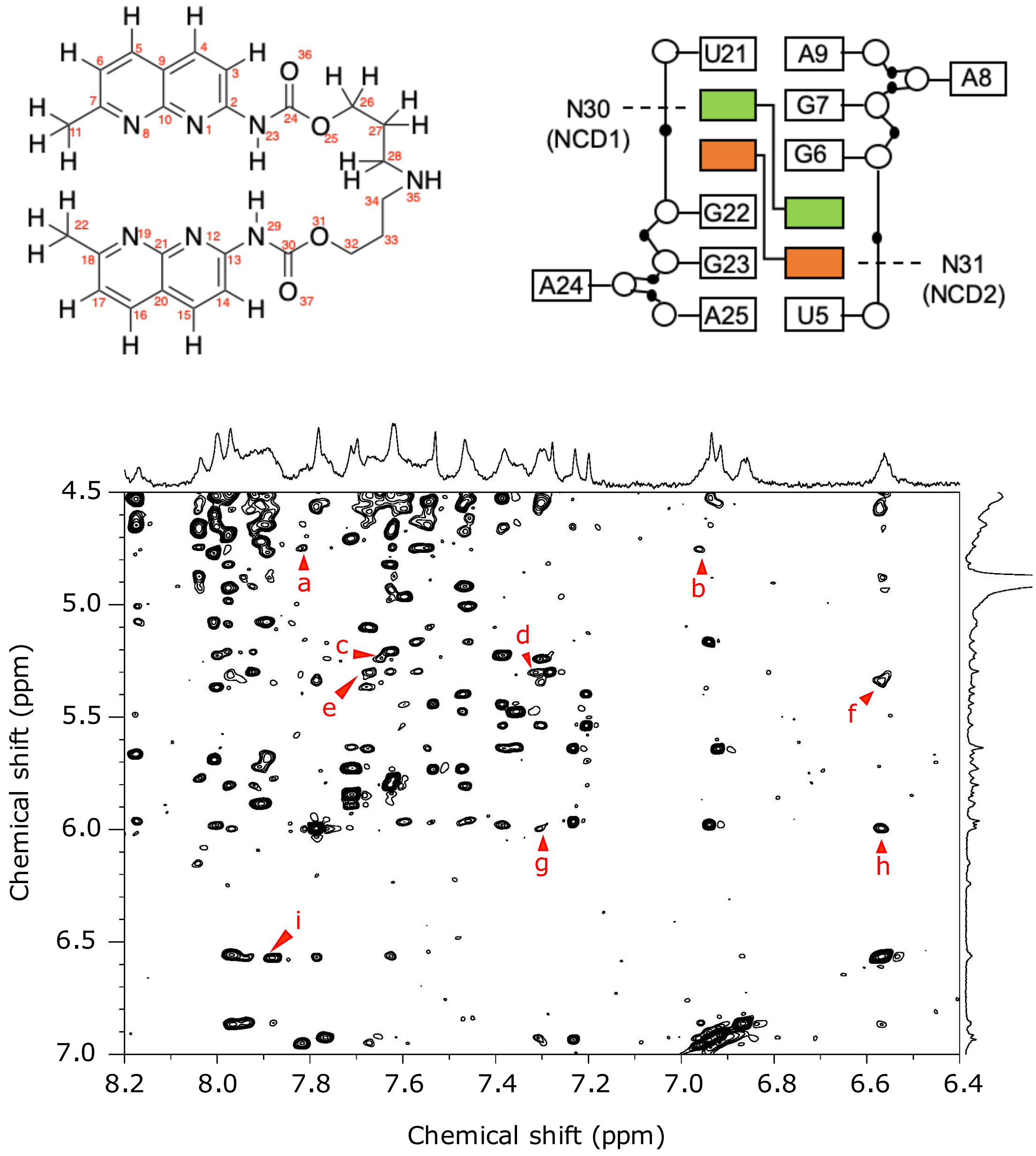

**Supplementary Figure 9**

Examples of intermolecular NOEs of SCA31 hpRNA and NCD (a) U5H5-N31H5, (b) U5H5-N31H6, (c) U21H1'-N30H4, (d) U5H1'-N31H3, (e) U5H1'-N31H4, (f) G22H2'-N31H14, G6H2'-N30H14, (g) G22H1'-N31H3, G6H1'-N30H3, (h) G22H1'-N31H14, G6H1'-N30H14, (i) N30H14-G6H8, N31H14-G22H8

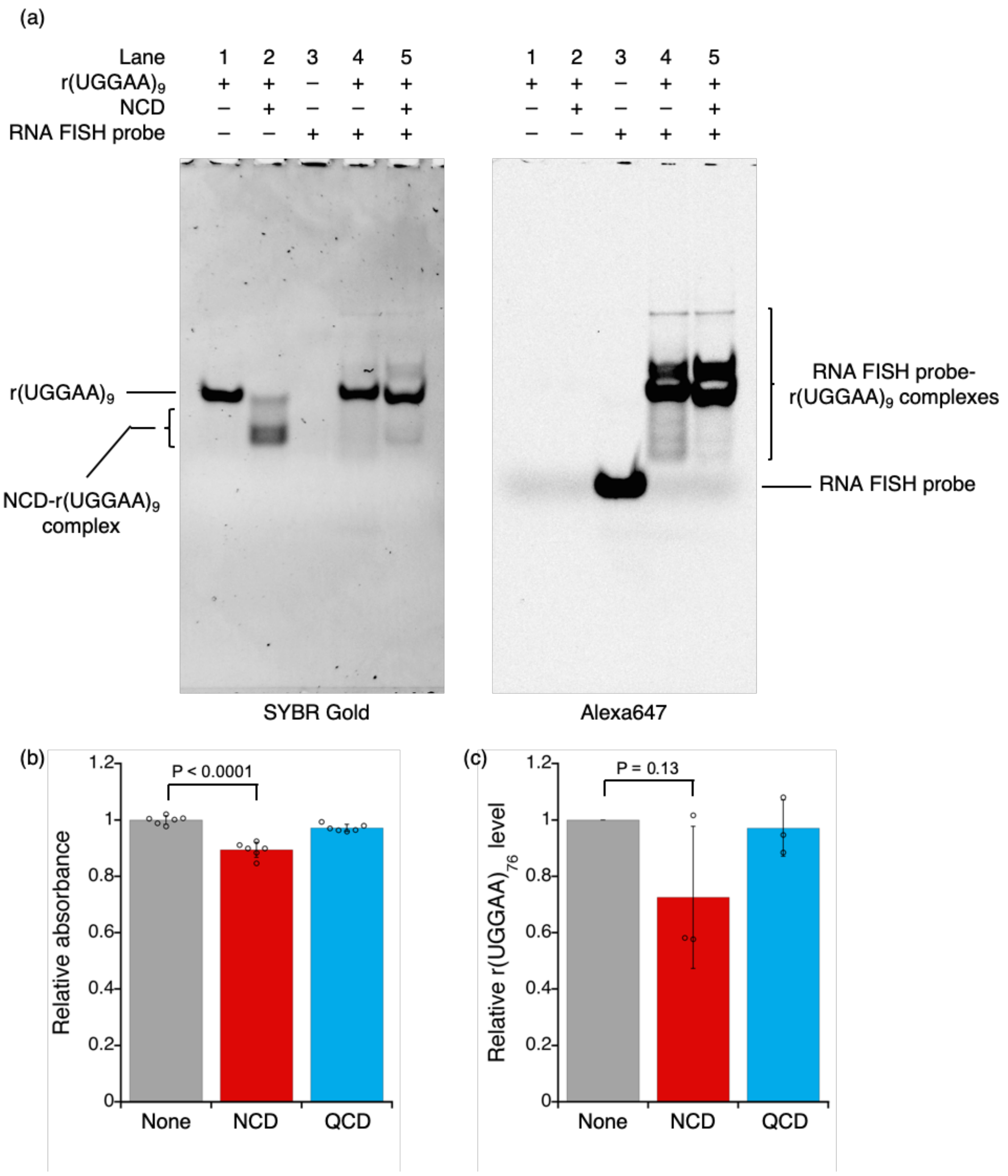

**Supplementary Figure 10**

(a) EMSA to confirm the interaction of r(UGGAA)$_9$ with RNA FISH probe in the absence and presence of NCD. r(UGGAA)$_9$ (200 nM) was incubated with RNA FISH probe (200 nM) in the absence and presence of NCD (2 μM) at 55 °C for 2 h. Bands were visualized by detecting SYBR Gold (left) and Alexa647 (right) with ImageQuant LAS 4000. RNA FISH probe: Alexa647-TTC5mCATT5mCCATTCCATT5mCCATTC5mCA, where underline is LNA and 5mC is 5-methylcytosine. (b) Cell viability assay and (c) Relative expression level of r(UGGAA)$_{76}$ in the absence and presence of ligands (5 μM). The *HPRT1* mRNA was used as an internal control. Data are shown as the mean ± SD (n = 5 for (b) and 3 for (c)); (Dunnett’s multiple comparison test).

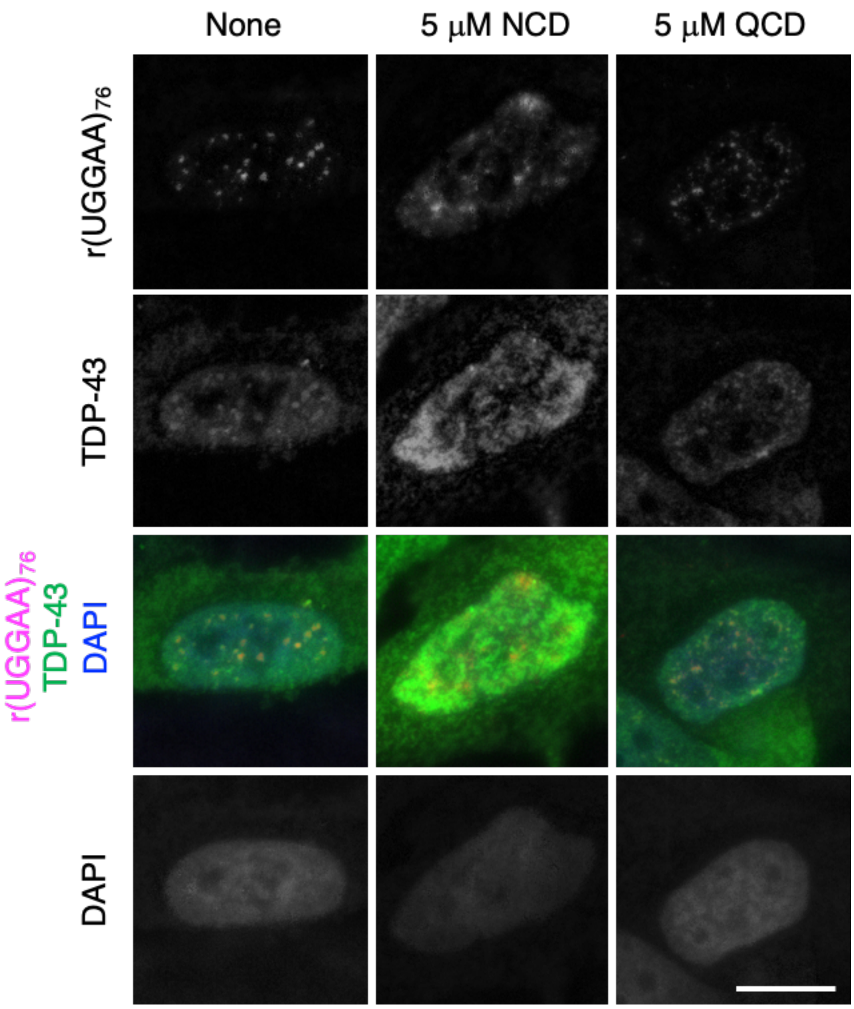

**Supplementary Figure 11**

RNA FISH and IF images of HeLa cells expressing r(UGGAA)$_{76}$ in the absence and the presence of NCD or QCD stained by r(UGGAA)$_{76}$-FISH and IF using anti-TDP-43 antibodies. Scale bar: 10 μm. In the control and QCD-treated HeLa cells, r(UGGAA)$_{76}$ colocalized with TDP-43, while the colocalization of r(UGGAA)$_{76}$ with TDP-43 was not observed in NCD-treated HeLa cells.

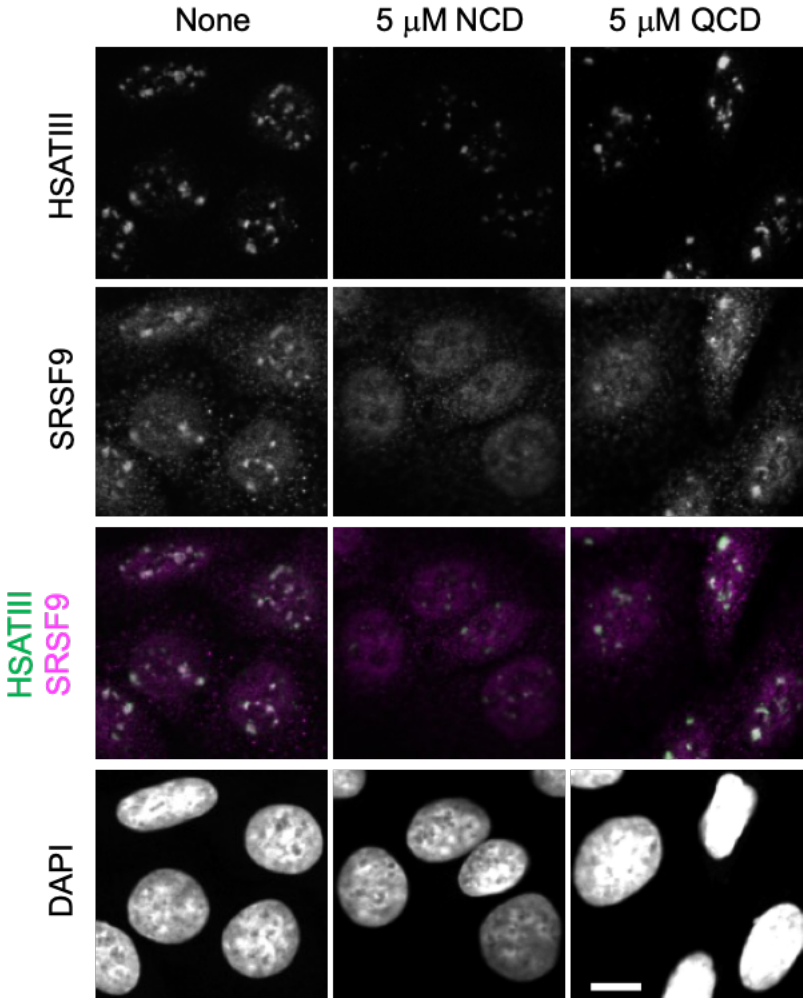

**Supplementary Figure 12**

RNA FISH and IF images of HeLa cells after thermal stress exposure in the absence and the presence of NCD or QCD stained by HSATIII-FISH and IF using anti-SRSF9 antibodies. Scale bar: 10 μm. SRSF9 colocalized with nSBs in the control and QCD-treated HeLa cells. In contrast, the decrease of nSBs and the diffusion of SRSF9 throughout nucleoplasm were observed in NCD-treated HeLa cells.

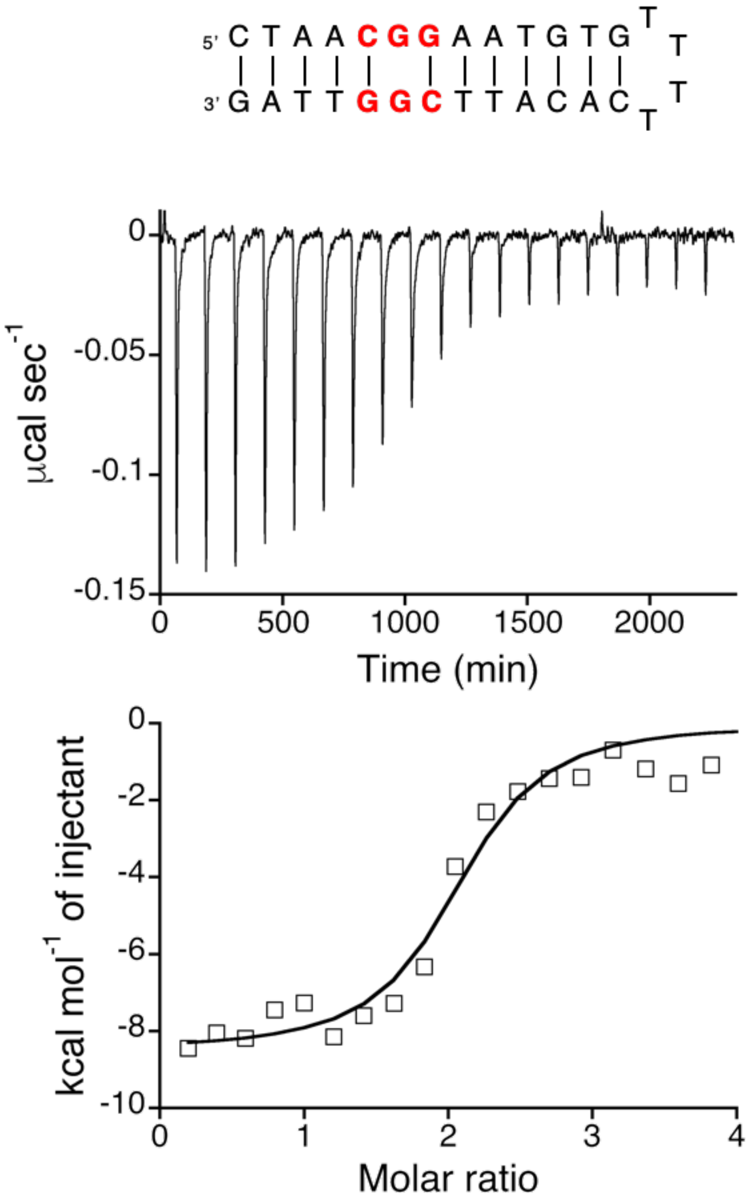

**Supplementary Figure 13**

ITC measurements for the binding of NCD to CGG/CGG-containing hairpin DNA.

**Supplementary Table 1. NMR constraints and structure statistics.**

| | |
|---|---|
| Number of experimental restraints | |
| Distance restraints | 192 |
| Intra-residue | 37 |
| Sequential | 53 |
| Long range | 21 |
| Medium range | 8 |
| Hydrogen bonding distance | 36 |
| Ambiguous | 37 |
| Dihedral restraints | 157 |
| Planarity | 14 |
| Heavy-atoms r.m.s. deviation (Å)[a] | |
| All including NCD | 1.966 ± 0.775 |
| All including NCD (pairwise) | 2.079 ± 0.359 |
| Backbone of RNA | 1.989 ± 0.787 |
| Backbone of RNA (pairwise) | 1.998 ± 0.396 |
| Backbone of RNA without triloop, A8, A24 | 1.553 ± 0.616 |
| Backbone of RNA without triloop, A8, A24 (pairwise) | 1.696 ± 0.321 |
| 5-7,9,21-23,25 with NCD | 1.069 ± 0.388 |
| 5-7,9,21-23,25 with NCD (pairwise) | 1.013 ± 0.145 |

[a]Averaged r.m.s.d. between an average structure and the 10 converged structures were calculated. The converged structures did not contain experimental distance violation of >0.5 Å or dihedral violation >5°.

### Supplementary Table 2. All-atom structure validation by MolProbity

| | | | | |
|---|---|---|---|---|
| 1 | | | | |
| All-Atom Contacts | Clashscore, all atoms: | 0 | | 100th percentile* (N=1784, all resolutions) |
| | Clashscore is the number of serious steric overlaps (> 0.4 Å) per 1000 atoms. | | | |
| Nucleic Acid Geometry | Probably wrong sugar puckers: | 2 | 6.90% | Goal: 0 |
| | Bad backbone conformations#: | 16 | 55.17% | Goal: <= 5% |
| | Bad bonds: | 0 / 690 | 0.00% | Goal: 0% |
| | Bad angles: | 1 / 1074 | 0.09% | Goal: <0.1% |
| Additional validations | Chiral volume outliers | 0/144 | | |
| | Waters with clashes | 0/0 | 0.00% | See UnDowser table for details |
| 2 | | | | |
| All-Atom Contacts | Clashscore, all atoms: | 0 | | 100th percentile* (N=1784, all resolutions) |
| | Clashscore is the number of serious steric overlaps (> 0.4 Å) per 1000 atoms. | | | |
| Nucleic Acid Geometry | Probably wrong sugar puckers: | 2 | 6.90% | Goal: 0 |
| | Bad backbone conformations#: | 16 | 55.17% | Goal: <= 5% |
| | Bad bonds: | 0 / 690 | 0.00% | Goal: 0% |
| | Bad angles: | 0 / 1074 | 0.00% | Goal: <0.1% |
| Additional validations | Chiral volume outliers | 0/144 | | |
| | Waters with clashes | 0/0 | 0.00% | See UnDowser table for detai |
| 3 | | | | |
| All-Atom Contacts | Clashscore, all atoms: | 0 | | 100th percentile* (N=1784, all resolutions) |
| | Clashscore is the number of serious steric overlaps (> 0.4 Å) per 1000 atoms. | | | |
| Nucleic Acid Geometry | Probably wrong sugar puckers: | 3 | 10.34% | Goal: 0 |
| | Bad backbone conformations#: | 15 | 51.72% | Goal: <= 5% |
| | Bad bonds: | 0 / 690 | 0.00% | Goal: 0% |
| | Bad angles: | 1 / 1074 | 0.09% | Goal: <0.1% |
| Additional validations | Chiral volume outliers | 0/144 | | |
| | Waters with clashes | 0/0 | 0.00% | See UnDowser table for details |
| 4 | | | | |
| All-Atom Contacts | Clashscore, all atoms: | 0 | | 100th percentile* (N=1784, all resolutions) |
| | Clashscore is the number of serious steric overlaps (> 0.4 Å) per 1000 atoms. | | | |
| Nucleic Acid Geometry | Probably wrong sugar puckers: | 1 | 3.45% | Goal: 0 |
| | Bad backbone conformations#: | 17 | 58.62% | Goal: <= 5% |
| | Bad bonds: | 0 / 690 | 0.00% | Goal: 0% |
| | Bad angles: | 0 / 1074 | 0.00% | Goal: <0.1% |
| Additional validations | Chiral volume outliers | 0/144 | | |
| | Waters with clashes | 0/0 | 0.00% | See UnDowser table for details |
| 5 | | | | |
| All-Atom Contacts | Clashscore, all atoms: | 0 | | 100th percentile* (N=1784, all resolutions) |
| | Clashscore is the number of serious steric overlaps (> 0.4 Å) per 1000 atoms. | | | |
| Nucleic Acid Geometry | Probably wrong sugar puckers: | 3 | 10.34% | Goal: 0 |
| | Bad backbone conformations#: | 14 | 48.28% | Goal: <= 5% |
| | Bad bonds: | 0 / 690 | 0.00% | Goal: 0% |
| | Bad angles: | 0 / 1074 | 0.00% | Goal: <0.1% |
| Additional validations | Chiral volume outliers | 0/144 | | |
| | Waters with clashes | 0/0 | 0.00% | See UnDowser table for details |
| 6 | | | | |
| All-Atom Contacts | Clashscore, all atoms: | 0 | | 100th percentile* (N=1784, all resolutions) |
| | Clashscore is the number of serious steric overlaps (> 0.4 Å) per 1000 atoms. | | | |
| Nucleic Acid Geometry | Probably wrong sugar puckers: | 1 | 3.45% | Goal: 0 |
| | Bad backbone conformations#: | 16 | 55.17% | Goal: <= 5% |
| | Bad bonds: | 0 / 690 | 0.00% | Goal: 0% |
| | Bad angles: | 0 / 1074 | 0.00% | Goal: <0.1% |
| Additional validations | Chiral volume outliers | 0/144 | | |
| | Waters with clashes | 0/0 | 0.00% | See UnDowser table for details |
| 7 | | | | |
| All-Atom Contacts | Clashscore, all atoms: | 0 | | 100th percentile* (N=1784, all resolutions) |
| | Clashscore is the number of serious steric overlaps (> 0.4 Å) per 1000 atoms. | | | |
| Nucleic Acid Geometry | Probably wrong sugar puckers: | 3 | 10.34% | Goal: 0 |
| | Bad backbone conformations#: | 14 | 48.28% | Goal: <= 5% |
| | Bad bonds: | 0 / 690 | 0.00% | Goal: 0% |
| | Bad angles: | 0 / 1074 | 0.00% | Goal: <0.1% |
| Additional validations | Chiral volume outliers | 0/144 | | |
| | Waters with clashes | 0/0 | 0.00% | See UnDowser table for details |

| | | | | |
|---|---|---|---|---|
| 8 | | | | |
| All-Atom Contacts | Clashscore, all atoms: | 0 | | 100th percentile* (N=1784, all resolutions) |
| | Clashscore is the number of serious steric overlaps (> 0.4 Å) per 1000 atoms. | | | |
| Nucleic Acid Geometry | Probably wrong sugar puckers: | 3 | 10.34% | Goal: 0 |
| | Bad backbone conformations#: | 15 | 51.72% | Goal: <= 5% |
| | Bad bonds: | 0 / 690 | 0.00% | Goal: 0% |
| | Bad angles: | 0 / 1074 | 0.00% | Goal: <0.1% |
| Additional validations | Chiral volume outliers | 0/144 | | |
| | Waters with clashes | 0/0 | 0.00% | See UnDowser table for details |
| 9 | | | | |
| All-Atom Contacts | Clashscore, all atoms: | 0 | | 100th percentile* (N=1784, all resolutions) |
| | Clashscore is the number of serious steric overlaps (> 0.4 Å) per 1000 atoms. | | | |
| Nucleic Acid Geometry | Probably wrong sugar puckers: | 2 | 6.90% | Goal: 0 |
| | Bad backbone conformations#: | 18 | 62.07% | Goal: <= 5% |
| | Bad bonds: | 0 / 690 | 0.00% | Goal: 0% |
| | Bad angles: | 1 / 1074 | 0.09% | Goal: <0.1% |
| Additional validations | Chiral volume outliers | 0/144 | | |
| | Waters with clashes | 0/0 | 0.00% | See UnDowser table for details |
| 10 | | | | |
| All-Atom Contacts | Clashscore, all atoms: | 0 | | 100th percentile* (N=1784, all resolutions) |
| | Clashscore is the number of serious steric overlaps (> 0.4 Å) per 1000 atoms. | | | |
| Nucleic Acid Geometry | Probably wrong sugar puckers: | 1 | 3.45% | Goal: 0 |
| | Bad backbone conformations#: | 15 | 51.72% | Goal: <= 5% |
| | Bad bonds: | 0 / 690 | 0.00% | Goal: 0% |
| | Bad angles: | 0 / 1074 | 0.00% | Goal: <0.1% |
| Additional validations | Chiral volume outliers | 0/144 | | |
| | Waters with clashes | 0/0 | 0.00% | See UnDowser table for details |
| average | | | | |
| All-Atom Contacts | Clashscore, all atoms: | | | 100th percentile* (N=1784, all resolutions) |
| | Clashscore is the number of serious steric overlaps (> 0.4 Å) per 1000 atoms. | | | |
| Nucleic Acid Geometry | Probably wrong sugar puckers: | 2.1 | | Goal: 0 |
| | Bad backbone conformations#: | 15.6 | | Goal: <= 5% |
| | Bad bonds: | | | Goal: 0% |
| | Bad angles: | | | Goal: <0.1% |
| Additional validations | Chiral volume outliers | | | |
| | Waters with clashes | | | See UnDowser table for details |

The all-atom structure validation by MolProbity indicated 15.6 bad backbone conformations per structure in the UGGAA/UGGAA pentad and GUUUC loop regions. Probably wrong sugar puckers were 2.1 per structure which were distributed mainly in the same regions. The conformation of the GUUUC loop region was not well defined due to the intrinsic flexibility. For the UGGAA/UGGAA pentad, it is suggested that the interaction with NCD induced the conformational change in this region. It is noted that the clashscore was zero and no bad bonds and three bad angles were found for the ten structures.

**Supplementary Table 3. Primer sequences used in RT-qPCR and semi-quantitative RT-PCR**

| | |
|---|---|
| HSATIII forward | 5’-TATGAATTCAATCAACCCGAGTGCAATCGAA-3’ |
| HSATIII reverse | 5’-TATGGATCCTTCCATTCCATTGCTGTACTCG-3’ |
| CLK1 exon 2 forward | 5’-ATGAGACACTCAAAGAGAACTTACTG-3’ |
| CLK1 intron 3 forward | 5’-TGCCGCCCACTTGACGTTTCCAG-3’ |
| CLK1 exon 6 reverse | 5’-TTACTGCTACATGTCTACCTCC-3’ |
| HSP-105 forward | 5'-ATGTTTCTGCACAGAAAGATGG-3' |
| HSP-105 reverse | 5’-TTTTGGGCTTTTTAGCTTCTGG-3’ |
| TNRC6a forward | 5’-GAATGTTACAAGACAAACGAAT-3’ |
| TNRC6a reverse | 5’-GTTGTGCTGCTGTGTTTCCA-3’ |
| r(UGGAA)$_{76}$ forward | 5’-CGAGCAGACATGATAAGATACATTGATG-3’ |
| r(UGGAA)$_{76}$ reverse | 5’-GCAATTGTTGTTGTTAACTTGTTTATTGC-3’ |
| GAPDH forward | 5’-ATGAGAAGTATGACAACAGCCTCAAGAT-3’ |
| GAPDH reverse | 5’-ATGAGTCCTTCCACGATACCAAAGTT-3’ |
| HPRT1 primer | Housekeeping Gene Primer Set (TAKARA 3790) |
